# Site-dependent transcriptomic signatures of endometriosis are conserved across hormonal states

**DOI:** 10.64898/2026.07.10.737856

**Authors:** Douglas E. Henze, George Crowley, Stephen R. Quake

## Abstract

Endometriosis, a chronic condition in which endometrial tissue grows at other sites in the body, produces lesions whose gene expression profiles vary by anatomical location and menstrual cycle stage. The extent to which these location-dependent transcriptional differences persist across hormonal states has remained unknown. Here, we compile a comprehensive single-cell RNA sequencing atlas of 672,051 cells across 112 donors from publicly available sources spanning the eutopic endometrium, menstrual effluent, ovaries, peritoneum, and ectopic endometrium from peritoneal and ovarian sites. By training machine learning classifiers on reference tissue signatures, we identify a subset of cells within ectopic lesions that retain a uterine transcriptomic identity, which we term “core” lesion cells, and which are distinguished from surrounding host tissue populations. We show that transcriptomic differences between core lesion cells at both ectopic sites are robust across all phases of the menstrual cycle and in patients receiving exogenous hormonal therapy, and validate these findings in independent datasets. Cell type- and tissue-specific gene signatures derived from these core populations are sufficient to classify disease status and lesion subtype in independent bulk tissue data across 168 patients. These findings establish that although the core lesion cells maintain elements of eutopic endometrium identity, they also contain information about the anatomical site of the lesion. This anatomic-specificity provides a potential framework for cycle-independent endometriosis diagnosis and subtyping.

## Main

Endometriosis, a chronic inflammatory disease affecting approximately 10% of reproductive-age women, is characterized by the growth of endometrial tissue outside of the uterus^1,2^. The clinical presentation is heterogeneous, ranging from dysmenorrhea (painful periods) and chronic pelvic pain to infertility, and diagnosis, which requires laparoscopic surgery with histological confirmation, is delayed by an average of 6-10 years^3,4^. Over this interval, disease can progress from minimal to severe stages as classified by the revised American Society for Reproductive Medicine (rASRM) system^5^. Beyond staging, endometriosis can be typed by lesion location, most commonly as ovarian endometriomas or peritoneal lesions, which are thought to represent distinct pathological entities with divergent structural architectures and potentially distinct pathogeneses^6^. However, no adequate non-invasive tools currently exist to distinguish one subtype from the other.

The growth and maintenance of ectopic lesions is driven in large part by estrogen signaling and disrupted progesterone responsiveness^7–9^, with additional menstrual hormones implicated in disease progression^10,11^. These hormonal dependencies have motivated the use of ovulation-blocking therapies as the primary medical treatment for endometriosis and have shaped the assumption that hormonal cycling is a major driver of transcriptomic heterogeneity within ectopic tissue. Indeed, the eutopic endometrium undergoes dramatic transcriptomic remodeling across the menstrual cycle^12,13^, and this responsiveness is at least partially preserved in ectopic lesions^6,14^. To investigate the cellular and molecular landscape of endometriosis across these hormonal states, single-cell RNA sequencing (scRNA-seq), a technique that measures gene activity in individual cells, has been applied to profile eutopic and ectopic tissues^14–17^. These studies have revealed transcriptomic heterogeneity across tissue sites and cell types, yet hormonal variation across the menstrual cycle and between treated and untreated patients remains deeply confounded with ectopic site, leaving it unclear which variable principally shapes the molecular identity of lesion subtypes.

Here, we integrate publicly available scRNA-seq datasets spanning the eutopic endometrium, menstrual effluent, ovaries, peritoneum, and ectopic endometrium from both peritoneal and ovarian sites to construct a comprehensive multi-tissue reference atlas. Using cell type-level classifiers trained on reference tissue signatures, we identify the core endometrial-like populations within each ectopic site and characterize how these populations differ from the surrounding host tissue microenvironment. We demonstrate that ectopic site, rather than hormonal state, is the primary axis of transcriptomic variation among core lesion mesenchymal lineage cells, indicating the robustness of potential RNA biomarkers across the menstrual cycle. Finally, we leverage the cell type- and tissue-specific signatures of core ectopic cells to stratify independent bulk tissue samples by disease status and lesion location, establishing a molecular framework for non-invasive endometriosis diagnosis that is independent of hormonal condition.

## Results

### Construction of a comprehensive multi-tissue reference atlas for endometriosis

To define cell types across endometriosis lesions in the context of their potential lesion sites, we assembled an integrated reference atlas comprising 672,051 cells from 112 donors spanning eutopic endometrium, menstrual effluent, ovaries, peritoneal biopsies, and ectopic endometrium from both peritoneal (EcP) and ovarian (EcO) sites (Figure 1a,b). The atlas was constructed by harmonizing publicly available single-cell RNA sequencing (scRNA-seq) datasets across these tissues. Specifically, to represent eutopic and menstruating endometrial tissue, we incorporated the Human Endometrial Cell Atlas (HECA)^17^, which provides harmonized metadata across publicly available eutopic endometrium samples, together with menstrual effluent profiles from Shih et al.^18^ that include healthy controls, symptomatic individuals, and endometriosis-positive individuals. To represent ovarian tissue, we integrated healthy ovarian datasets from Jones et al.^19^, Tabula Sapiens 2.0^20^, and Fonseca et al^15^. To include ectopic tissue and enable direct comparisons between both peritoneal and ovarian ectopic sites without introducing sampling bias, we restricted inclusion to studies that profiled both EcP and EcO lesions. This included Fonseca et al. and Tan et al.^16^. To incorporate a peritoneal tissue reference, we also included the EcP-adjacent samples from Tan et al., and the unaffected ovary samples from Fonseca et al. as an additional ovary reference. Although these datasets originated from different laboratories and collection time points, they exhibited comparable read depth and gene detection rates (Supplementary Figure 1a,b), supporting their suitability for joint integration.

**Figure 1:**
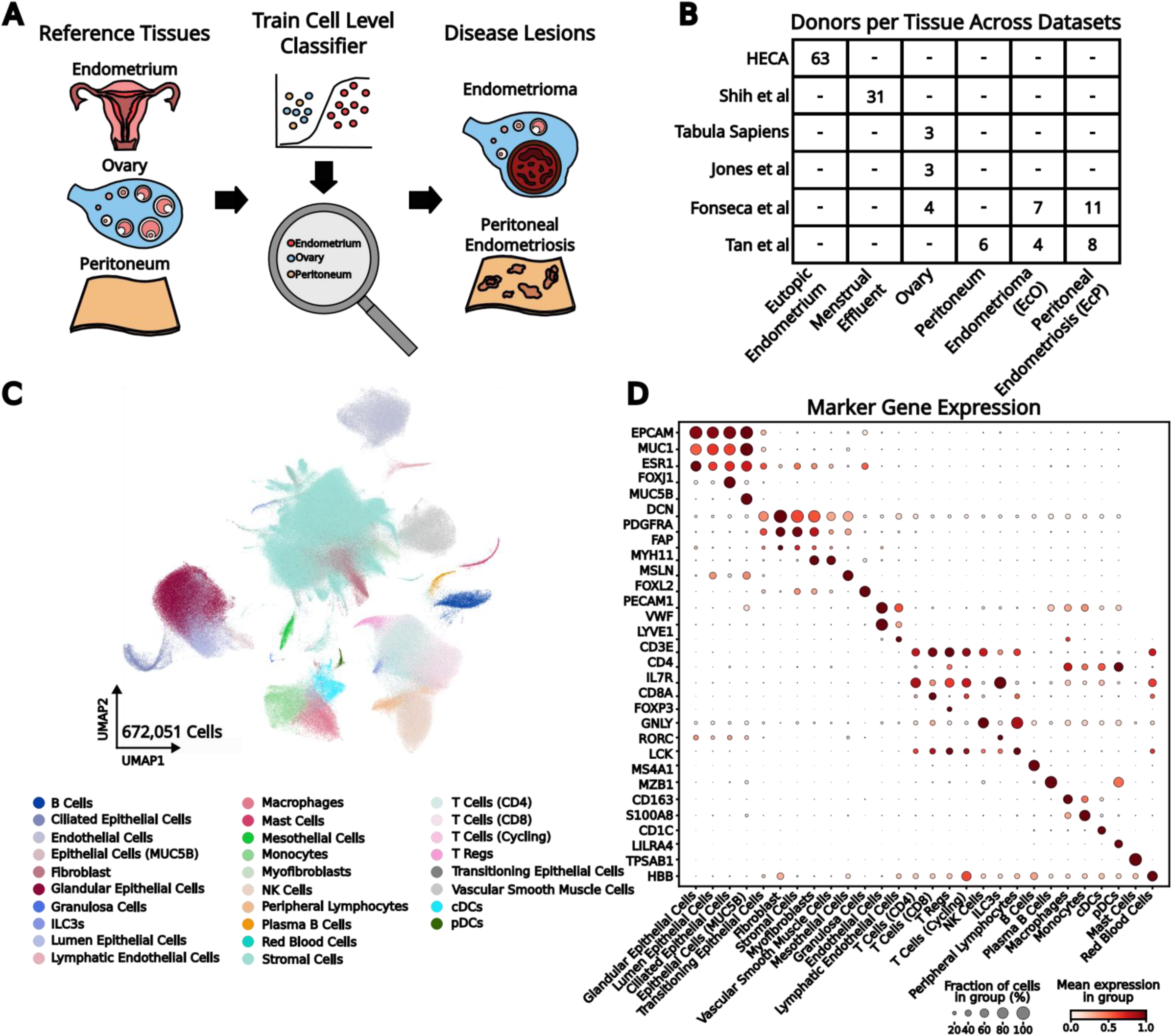
Integrated cell atlas across endometriosis related tissues. **A,** Schematic overview of the data analysis pipeline, building references and training classifiers on reference tissue and applying them to lesions. **B,** Donor distribution, number of unique donors for every tissue-dataset pair. **C,** UMAP plot across the different tissues colored by broad cell type. **D,** Marker gene expression across the broad cell types.

To establish unified cell identities across these diverse tissues, we performed large language model (LLM)-based *de novo* cell type annotation^21^, generating broad cell class labels across all integrated datasets (Figure 1c; Methods). The resulting LLM annotations mapped closely to the expert-curated annotations in the HECA (Supplementary Figure 1c) and aligned well with canonical marker gene expression for each cell type (Figure 1d), validating the unified labels. The most notable discrepancies between the LLM-based and HECA annotations involved T cell subpopulations and red blood cells; the LLM-guided approach classified a greater proportion of cells as blood cells, consistent with a conservative interpretation of the strong hemoglobin signal observed broadly across the atlas in previous studies (Figure 1d). Beyond reproducing known annotations, integration coupled with LLM-based labeling highlighted additional populations and transitional states not emphasized in the source studies. These included myofibroblast-like cells in the eutopic endometrium (Figure 1d, Supplementary Figure 1c) and transitional epithelial cell states co-expressing canonical epithelial and stromal markers. Additionally, we identified a mesothelial cell population, which was most abundant in the ectopic peritoneal tissue (Figure 1c, Supplementary Figure 1d).

Taken together, atlas integration substantially reduced batch effects across tissues, datasets, and disease states (Supplementary Figure 1d-f) while yielding unified cell type annotations spanning all sampled compartments. This comprehensive multi-tissue reference atlas provides a foundation for defining tissue-specific transcriptomic signatures and for deconvolving cell type-level signals in endometriosis.

### Somatic mutations link ectopic lesions to eutopic endometrium

The prevailing hypothesis for the pathogenesis of endometriosis is that retrograde menstruation leads to endometrial tissue implanting within the peritoneal space^22^. If ectopic lesions originate from displaced eutopic endometrium, then both tissues should share a common mutational history. To assess this, we identified 1,064 high-confidence somatic variants across paired eutopic (300 variants) and ectopic (764 variants) endometrium samples in the Fonseca et al. dataset (Methods) (Supplementary Figure 2a). We first compared the mutational processes operating in each tissue by analyzing their mutation spectra. The 6-class mutation spectrum revealed similar proportional contributions in both tissues (Supplementary Figure 2a), and the 96-class trinucleotide spectrum confirmed broad similarity between the two sites (r = 0.87) (Supplementary Figure 2b). Both spectra exhibited a prominent SBS5 signature, characterized by frequent C>T and T>C mutations, consistent with prior observations across eutopic and ectopic endometrium^23,24^.

Having established that both tissues undergo similar mutagenic processes, we next asked whether the identified somatic mutations were subject to selection. We computed the ratio of nonsynonymous to synonymous mutations (dN/dS) and evaluated whether the observed ratio exceeded random expectations by generating null distributions across 100,000 permutations. In each permutation, an alternative base was randomly assigned, weighted by the observed trinucleotide spectrum, at the sequence-specific context of each coding variant position (Methods). The observed dN/dS was indistinguishable from the null distribution (Supplementary Figure 2c), indicating that somatic mutations accumulate without detectable genome-wide selection for or against protein-altering changes. Together with the shared mutagenic signatures, this suggests that both tissues harbor a set of passenger mutations amenable to lineage tracing.

We therefore asked whether any somatic variants were shared between paired eutopic and ectopic samples, and identified sets of shared mutations in two patients (Supplementary Figure 2d). To assess whether these shared variants reflected clonal relationships, we analyzed the clonal cell fraction, defined as the proportion of cells confidently carrying each mutation (Methods). In both patients, the shared mutations were present in comparably small fractions of cells at both sites (Supplementary Figure 2e), consistent with an oligoclonal structure in which the ectopic lesion does not arise from a single massively expanded clone. To determine whether any clonal expansion nonetheless occurred at ectopic sites, we compared variant allele frequencies (VAFs) of all called mutations across paired eutopic and ectopic samples in all four patients (Supplementary Figure 2f). Across four of five paired samples, ectopic sites exhibited modestly elevated VAFs, suggesting a degree of clonal expansion following the initial seeding event.

Finally, to understand whether somatic mutations accumulate differentially across cell types, we mapped called variants back to their cell type of origin (Supplementary Figure 2g). Consistent with previous results^25^, the stromal and fibroblast fraction carry variant alleles at a relatively abundant rate. This result is not consistent with prior bulk observations that epithelial cells harbor the most mutations, likely owing to lower cell numbers that led to reduced sequencing depth in the bulk analysis. Notably, the distribution of mutations between eutopic and ectopic endometrium varied substantially across cell types, motivating a cell-type-resolved analysis of the ectopic lesion.

### Cell type-level classifiers distinguish endometrial-like from host tissue cells within ectopic lesions

During atlas integration, we observed that certain ectopic endometrium stromal cell populations co-clustered with ovarian stromal cells (Supplementary Figure 1d), raising the question of whether an underlying transcriptomic signature for each reference tissue could be leveraged to classify cell types within the ectopic endometrium as endometrial derived or host-tissue derived with high confidence (Figure 1a). To test this, we trained logistic regression classifiers on differentially expressed, highly cell type-specific genes to distinguish endometrial (eutopic/menstrual effluent) from non-endometrial reference (ovary/peritoneum) cell types (Methods). The classifiers achieved strong performance across the most abundant cell types (Figure 2a) and accurately assigned the majority of cells as endometrial or non-endometrial in origin (Supplementary Figure 3a), with a pooled false discovery rate of 1.5% across all cell types. Notably, the feature genes driving the majority of our cell type-level classifiers were enriched for endometrium-specific genes as defined by RNA expression in the Human Protein Atlas (HPA)^26^ (Supplementary Figure 3b), further validating the biological relevance of the discriminative signal.

**Figure 2:**
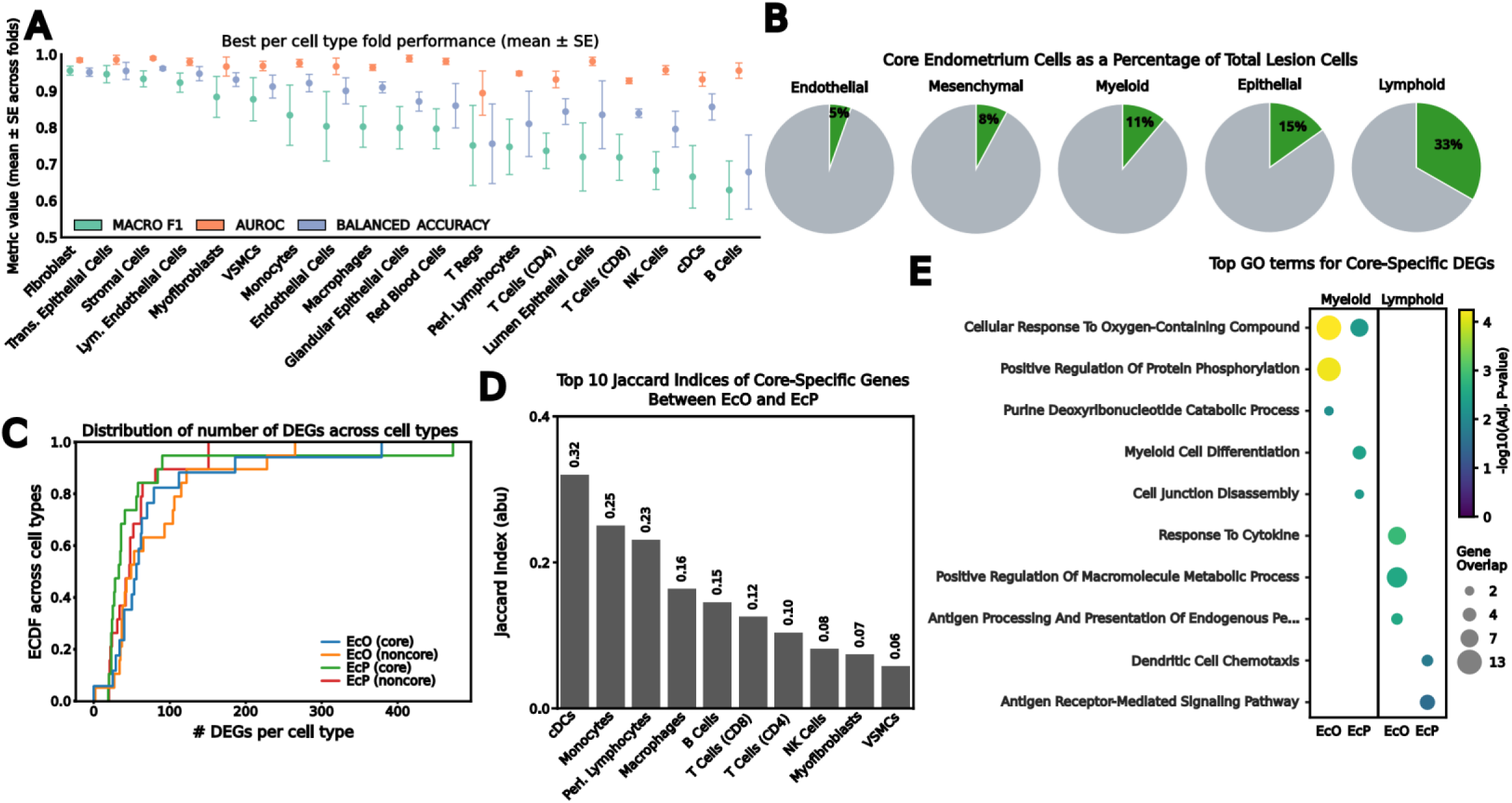
Tissue-of-origin classifiers reveal heterogeneous endometrial-like and host tissue populations within ectopic lesions. **A,** Grouped bar plot showing the relative performance amongst each of the cell type-specific classifiers. Sorted by macro F1. **B,** Percentage of lesion cells for each lineage which mapped to being endometrium-like (green) or not (gray). **C,** Empirical cumulative distribution function to show the relative percent of differentially expressed genes from the eutopic endometrium; lines are colored by site and core or non-core category. **D,** Ranked Jaccard Indices between the core-specific differentially expressed genes for each of the cell types across the endometrioma and peritoneal endometriosis samples **E,** Gene ontology biological processes terms for the differentially expressed genes from the eutopic endometrium for myeloid and lymphoid cells within each of the sites.

Given that endometriosis is defined by the growth of endometrial-like tissue outside the uterus, we next sought to determine whether cells within resected ectopic sites exhibited an endometrial transcriptomic identity. Applying our classifiers across the entire dataset (Supplementary Figure 3c), we found that the majority of cells at ectopic sites were transcriptomically more similar to their host tissue of residence than to the endometrium. The proportion of lesion cells classified as endometrial-like, hereafter referred to as “core” lesion cells, varied by lineage: endothelial cells showed the lowest fraction (5%), followed by mesenchymal (8%), myeloid (11%), epithelial (15%), and lymphoid (33%) lineage cells. These modest proportions are consistent with prior observations that stromal cells of endometrial origin constitute a minority of endometrioma tissue^14^ and that immune populations within lesions represent an admixture of host- and endometrium-derived cells^27^. Comparing across ectopic sites, we observed broadly comparable rates of endometrial-like classification, with the most notable divergence occurring among immune lineages, where EcO myeloid cells and EcP lymphoid cells were more frequently classified as endometrial-like (Supplementary Figure 3 d,e). The relative enrichment of endometrial-like myeloid cells in EcO is consistent with previous observations that the predominant macrophage population in endometriomas exhibits a tissue-resident phenotype that is also enriched in the eutopic endometrium of patients with endometriosis^16^.

To test whether core populations faithfully recapitulate the eutopic endometrial microenvironment, we performed differential cell-cell interaction analysis^28^ comparing the eutopic endometrium across all endometriosis patients against the core cells of both the EcO and the EcP sites (Methods). In the EcO, core cells retained conserved interactions among mesenchymal populations relative to non-core cells (Supplementary Figure 4a), suggesting preservation of structural cell communication networks. The EcP maintained this mesenchymal interaction, but it also displayed a greater number of enriched core interactions involving epithelial populations, more closely resembling the eutopic endometrium than the EcO (Supplementary Figure 4b). This divergence likely reflects the altered structural dynamics between EcO and EcP, as well as the reduced prevalence of epithelial tissue within endometrioma samples^14–16^. Consistent with prior reports, core EcP cells also exhibited well-conserved epithelial-to-stromal crosstalk^29^.

To directly assess transcriptomic similarity to eutopic endometrium, we performed differential expression analysis between diseased eutopic endometrium and both core and non-core populations within each ectopic site separately (Supplementary Figure 5a,b; Methods). Across cell types, core populations consistently exhibited fewer differentially expressed genes (DEGs) relative to diseased eutopic endometrium than non-core populations at the same site (Figure 2c), suggesting that core cells are transcriptomically closer to endometrial tissue. We also observed a higher DEG burden per cell type in EcO than in EcP core cells, suggesting that endometrial-like populations in endometrioma may be more divergent from diseased eutopic endometrium than those in peritoneal lesions.

To interpret the biological basis of these differences, we partitioned DEGs for each cell type into core-specific, non-core-specific, and shared gene programs between EcO and EcP. We focus on macrophages, which exhibited amongst the largest DEG burdens at both sites (Supplementary Figure 5a,b), to illustrate the functional differences between ectopic sites. In EcO, non-core-specific macrophage DEGs were enriched for cell migration pathways (Supplementary Figure 5c), consistent with evidence that macrophages can traffic from host tissue into endometrial lesions^30^. The core-specific and shared programs in EcO macrophages highlighted cytokine response and positive regulation of smooth muscle proliferation, with the core-specific program additionally including activation pathways (Supplementary Figure 5d,e), aligning with the inflammatory milieu of endometriosis and macrophage contributions to lesion vascularization^31,32^. In EcP, non-core-specific genes reflected a macrophage expansion response similar to that in EcO, but with the addition of local proliferation pathways rather than solely migration, and with a stronger extracellular matrix organizing component, consistent with prior results^33^ (Supplementary Figure 5f). EcP shared and core-specific macrophage signatures showed enrichment for golgi lumen acidification and positive regulation of cell death, respectively (Supplementary Figure 5g,h), the latter raising the possibility of macrophage-associated dysfunction within the lesions.

Having established that core-specific DEGs capture distinct biology characterizing ectopic cells relative to eutopic tissue, we next asked how conserved these transcriptomic programs were between lesion sites. Strikingly, Jaccard indices between EcO and EcP core-specific gene sets were relatively low across individual cell types (Figure 2d), with the greatest site-conserved changes occurring within immune populations. Grouping core-specific DEGs by immune lineage revealed that the majority of enriched programs were site- and lineage-specific. Specifically, lymphoid DEG pathways reflected chemotaxis in EcP and response to cytokines in EcO, while myeloid pathways reflected differentiation programs in EcP and catabolic processes in EcO (Figure 2f). These site-dependent transcriptomic signatures suggest that the local tissue environment makes a substantial contribution to the molecular identity of endometrial-like cells.

### Menstrual cycle stage and exogenous hormones modulate ectopic endometrium in a cell type-specific manner

Fluctuating estrogen and progesterone levels across the menstrual cycle are widely thought to modulate the function of the endometrial lesion-resident cell types, analogous to hormone-responsive programs in the eutopic endometrium^12,17^. We asked whether these changes within the ectopic endometrium could explain the varying degrees of pain experienced throughout the menstrual cycle by endometriosis patients. These patients are known to experience primary dysmenorrhea, which is an acute symptom limited to the perimenstrual window and driven by myometrial hypercontractility, vasoconstriction and ischemic pain^34^. In addition to this primary pain, endometriosis patients experience a secondary dysmenorrhea, which can begin in the secretory phase of the menstrual cycle^35,36^, indicative of a pain mechanism which is entirely separate from the hypercontractility in primary dysmenorrhea^37^. We first quantified the number of genes varying along the menstrual cycle for each cell type with respect to its compartment of origin (Methods). This revealed substantial differences in hormone responsiveness between the eutopic endometrium and the ectopic sites for both immune and stromal cell compartments (Supplementary Figure 6a).

Given that the menstrual cycle comprises two distinct phases (proliferative and secretory) associated with different levels of endometriosis-related pain, and that exogenous hormones constitute an additional hormonal state, we next sought to identify phase-dependent expression patterns within each cell type. We calculated mean expression profiles of gene–cell type–tissue triplets across the three hormonal strata and clustered these profiles into six groups with similar expression dynamics (Supplementary Figure 6b; Methods). To identify the underlying gene programs, we annotated the clusters using gene ontology biological processes (Methods), which revealed correspondence with established endometrial biology. Because endometriosis-associated pain is initiated in the secretory phase^35^, we focused on the three clusters with expression peaks or troughs in this interval. Cluster 3, which peaks in the secretory phase, was characterized by the response to zinc ion, an element known to fluctuate throughout the menstrual cycle and accumulate during the secretory phase^38^. Conversely, Cluster 1, which troughs in the secretory stage, was enriched for extracellular matrix assembly, consistent with post-menstrual stromal regeneration and matrix deposition^39^, while Cluster 6, which also reached its lowest expression in the secretory phase, was defined by antigen presentation and alpha beta T cell activation with simultaneous suppression of NK cell-mediated cytotoxicity, inconsistent with prior observations of immune tolerance in the secretory state^40,41^. This failure to adopt the tolerogenic immune state that normally characterizes the secretory phase^41^ may reflect a blunted progesterone response, a hallmark of endometriosis driven by loss of the stimulatory PR-B isoform^42,43^.

We partitioned the cycle-responsive genes in the peaking and troughing groups by their tissue and cell type of origin (Supplementary Figure 6c, d). The greatest number of cycle-responsive genes occurred in mesenchymal populations, including stromal cells, fibroblasts, and vascular smooth muscle cells. Notably, across all cell types, these genes exhibited site-specific signatures, indicating that eutopic and ectopic endometrium diverge in their hormonal response.

To characterize the functional implications of secretory-phase dynamics restricted to ectopic tissue, we took the union of EcO and EcP genes across cell types. The secretory-peak programs were most strongly enriched for macrophage immune response and fibroblast ECM development (Supplementary Figure 6e), consistent with the heightened inflammatory and fibrotic states reported in endometriotic lesions^33,44^. Programs troughing in the secretory state further reinforced an inflammatory signature, with macrophages displaying elevated cytokine production (Supplementary Figure 6f). Unexpectedly, NK cells exhibited increased cytotoxicity, diverging from their canonical behavior during the secretory phase^45^ (Supplementary Figure 6f), reinforcing the position that the ectopic immune compartment becomes inflammatory at the initiation of secondary dysmenorrhea^37^. In contrast, monocytes contributed an anti-inflammatory profile, largely aligned with prior evidence that macrophage ontogeny and recruitment status give rise to functionally divergent populations within endometriotic lesions^27,46^. The convergence of these inflammatory and cytotoxic profiles within fibrotic lesions provides a molecular framework for the secondary dysmenorrhea of endometriosis that is mechanistically distinct from primary. In place of an acute myometrial contraction, the lesion’s immune cells sustain a hormonally driven inflammatory state linking hormonal fluctuations to the modulation of pain across the menstrual cycle in endometriosis patients.

### Ectopic site modulates cell type-specific transcriptomic programs independently of hormonal state

While our analysis shows that ectopic endometrial cells are hormone responsive, it also revealed pronounced, site-specific transcriptomic divergences from the eutopic endometrium among core ectopic cell types. We therefore asked which effect was larger: whether the transcriptomic variation in the ectopic endometrium primarily reflects changes in hormonal state or whether the ectopic site itself imposes an independent and dominant effect on cell state (Figure 3a).

**Figure 3:**
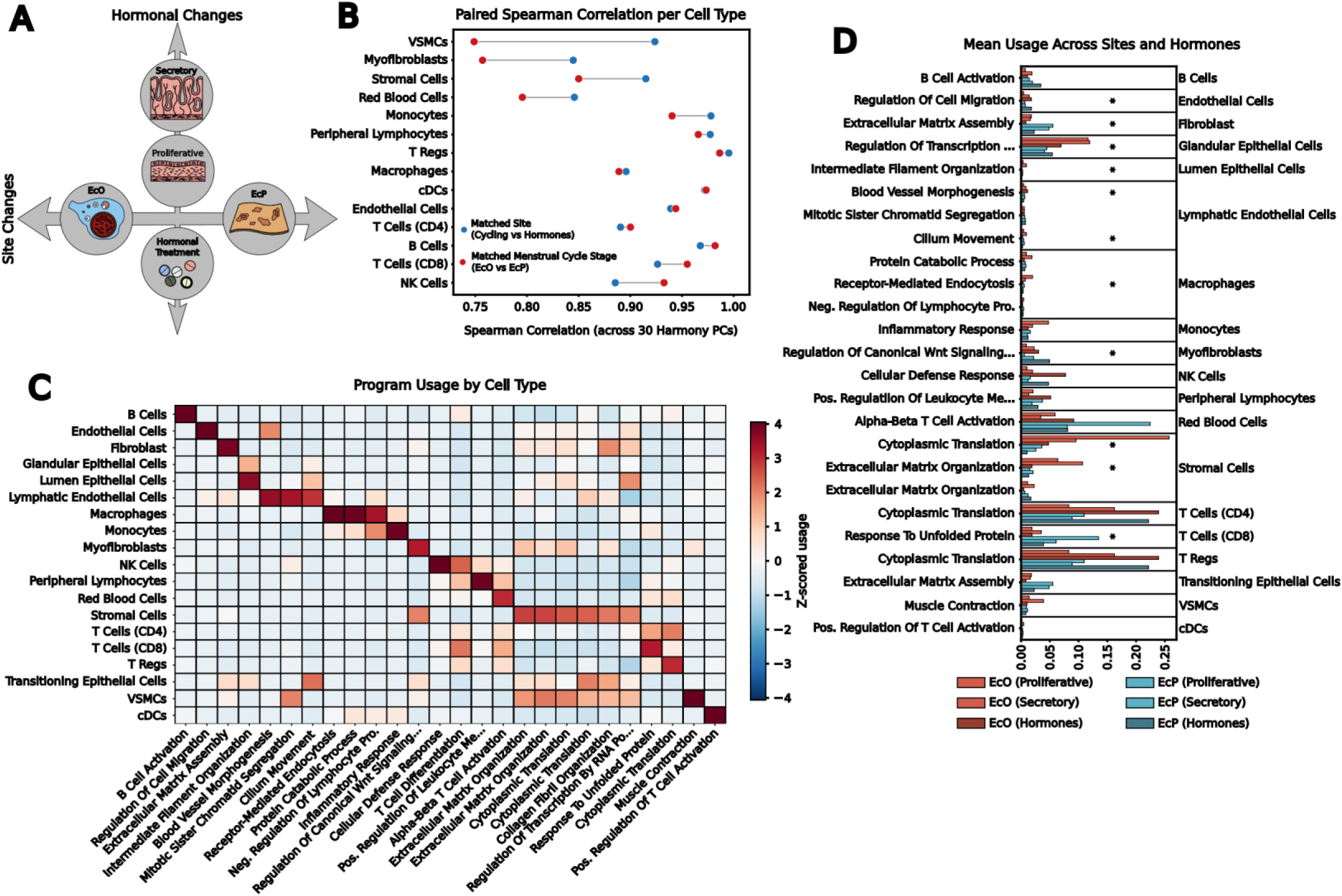
Transcriptomic programs are more associated with site than hormonal status. **A,** Schematic image of the primary axes of lesion cell variation being hormonal or spatial. **B,** Dumbbell plot showing the spearman correlation across sites in matched hormone cases or across hormones in matched site cases, with line meaning perfect agreement. **C,** Diagonalized heatmap showing the z-scored usage of different NMF modules across cell types. **D,** Bar plot showing the mean usage for each site-hormone pair across all cell types. * represents a statistically significant deviation between sites via a linear mixed model.

To disentangle these two variables, we computed Spearman correlations between the mean batch-corrected principal component coordinates of each core cell type under two conditions: first, between EcO and EcP cells in matched hormonal states, and second, between naturally cycling cells and those from patients receiving exogenous hormonal treatment at matched sites. A greater number of cell types exhibited higher correlation between different hormonal states than between different sites (Figure 3b), indicating that ectopic site exerts a stronger influence on transcriptomic identity than hormonal status. This effect was not uniform across compartments. The gap between site and hormonal correlations was widest for mesenchymal populations, including stromal cells, myofibroblasts, and vascular smooth muscle cells, whereas immune cell types were comparatively stable across both perturbations, with slightly higher correlation between sites than hormonal states (Figure 3b). This finding is notable given the longstanding emphasis on interpreting hormonal cycling as a primary driver of transcriptomic variation in ectopic endometrial tissue. To systematically characterize this site-dependent heterogeneity, we identified co-expressed gene modules within the same cell types across all donors, ectopic sites, and hormonal states (Methods). This analysis yielded thirty-three gene expression modules, twenty-six of which were significantly enriched for at least one Gene Ontology Biological Process term and were retained for annotation (Figure 3c; Supplementary Data Table 1; Methods). These modules were variably used across cell types, with each module showing its highest mean usage in a single cell type (Figure 3c).

We next restricted analysis to modules whose usage was specific to a cell type, which yielded twenty-two modules, and three of the cell types had no specific module and were assigned the most highly used annotated module, giving twenty-five cell type-module pairs spanning twenty-three unique modules (Supplementary Data Table 2; Methods). We then tested whether module usage differed between EcO and EcP independently of hormonal state (Methods). Eleven of the twenty-five cell type-module pairs showed significant site-dependent deviation, and these were predominantly associated with structural and endothelial populations, with only two immune-associated program reaching significance, suggesting that the structurally important cell types within the lesions are what drive site dependence (Figure 3d; Supplementary Data Table 3). Gene set enrichment analysis of these site-dependent modules converged on several recurring functional axes. Endothelial cells preferentially engaged cell migration programs involving *CDH5*, *CLDN5*, and *ROBO4* (Supplementary Data Table 1), consistent with established roles for endothelial angiogenesis^47^ in endometriosis. Lymphatic endothelial cells showed increased enrichment of blood vessel morphogenesis programs in EcP across both the Secretory and Exogenous Hormones states, in line with reports of increased lymphatic vessel density in peritoneal endometriosis^48^.

Epithelial lineage cells diverged between ectopic sites through intermediate filament assembly programs involving *KRT19*, *KRT17*, and *KRT16* (Supplementary Data Table 1), genes implicated in recognized hallmarks of endometriosis such as tissue remodeling and epithelial-mesenchymal transition^49^. Among mesenchymal cell types, extracellular matrix (ECM) organization programs included genes such as *GAS6* (Supplementary Data Table 1), a known regulator of endometriosis-related fibrosis, ECM remodeling, and immunosuppression in other disease contexts^50,51^. Myofibroblasts showed deviations in pathways involving *IGFBP1*, *IGFBP2*, and *IGFBP4* (Supplementary Data Table 1), which exhibit differential expression across lesion sites^52^ and regulate fibrosis^53^ and stromal cell proliferation^54^ in endometriosis. Stromal populations displayed increased usage of programs related to cytoplasmic translation and ECM organization, reflecting the extensive fibrosis and remodeling characteristic of different lesion sites^6,14^. Among the immune cell programs with significant site dependency, CD8 T cells were related to the unfolded protein response, encompassing heat shock proteins indicative of the oxidative stress environment in endometriosis^55^. Collectively, these divergences reveal a previously unappreciated site dependency of endometriosis-associated transcriptomic programs spanning structural, epithelial, and immune cell compartments.

Although these programs demonstrated site-specific enrichment, their usage varied across hormonal states, prompting us to ask whether the site-dependent differences persisted across different hormonal milieus. Remarkably, the EcO-to-EcP program deviations across all cell types were conserved regardless of whether individuals were receiving exogenous hormones or menstrual cycle stage (Supplementary Figure 7a), indicating that the observed site-specific differences were independent of hormonal state. To validate this finding, we performed differential expression analysis between EcO and EcP mesenchymal lineage core lesion cells from donors receiving exogenous hormones in the Tan et al. dataset and applied hierarchical clustering of the top differentially expressed genes to the same cell types across a range of hormonal states in the entirely held-out Fonseca et al. dataset. Strikingly, EcO and EcP cells clustered apart from one another regardless of hormonal state or cell type (Supplementary Figure 7b), demonstrating that site-driven transcriptomic differences are robust across the full spectrum of hormonal conditions. The genes driving this separation carry mechanistic relevance in endometriosis. *TNXB*, *MME*, and *RHOB*^56–58^ have each been implicated in disease pathogenesis through ECM remodeling or epithelial-mesenchymal transition, while *TCF21* and *IL32* are recognized regulators of the inflammatory phenotype in endometriosis^59,60^. *ID4* has been identified through genome-wide association studies (GWASs) as having an increased association with severe relative to mild endometriosis^61^, raising the possibility that these site-dependent signatures may also capture variation in disease severity independent of rASRM staging.

### Cell type lineages exhibit transcriptomic responses varying with endometriosis severity

The identification of a severity-associated gene within our site-dependent signatures prompted us to ask more broadly whether cell type lineages in ectopic lesions exhibit differential transcriptomic responses to increasing disease severity, and whether the differentially expressed genes shared across severity stages differ by lesion site. To address this, we performed differential expression analysis comparing control healthy reference tissue against non-core lesion cells and eutopic endometrium against core lesion cells for each lineage, then assessed whether the resulting effect sizes scaled with disease severity (Methods). Comparing these effect sizes between mild and severe endometriosis across lineages, we observed differential directionality: vascular and epithelial cell populations displayed decreasing effect sizes with greater severity, whereas immune populations showed increasing effect sizes (Supplementary Figure 8a), suggesting lineage-dependent transcriptomic remodeling as disease progresses.

Given these divergent lineage responses, we next examined whether the intersections between differentially expressed genes in mild and severe endometriosis at each lesion site were distinct. The EcP sites exhibited a more conserved pan severity signature than the EcO sites (Supplementary Figure 8b, c), indicating less overall transcriptomic difference associated with the EcP. Among the shared differentially expressed genes, both sites displayed similar proportions of cell type-specific and tissue-specific signatures, comprising 30 cell type-specific genes identified as cell type-specific in Tabula Sapiens^62^ alongside 14 genes with endometrium, ovary, or peritoneum specificity as annotated by the Human Protein Atlas^63^ (Supplementary Figure 8b,c; Methods). This co-occurrence of both cell type- and tissue-specific genes within severity-shared signatures raised the question of whether such signatures carry diagnostic utility beyond our single-cell framework.

### Cell type- and tissue-specific gene signatures are sufficient to stratify endometriosis subtypes

To assess the diagnostic potential of the cell type- and tissue-specific signatures identified through the severity analysis, we turned to independent bulk tissue microarray data spanning eutopic endometrium and peritoneum from both endometriosis patients and control individuals, as well as bulk lesion samples from deep infiltrating lesions, superficial peritoneal lesions, and endometriomas^64^ (Figure 4a; Supplementary Figure 8a,b). Projecting the bulk expression of both the cell type- and tissue-gene sets into PCA space revealed separation among tissue types (Supplementary Figure 9a,b), and we hypothesized that a combined signature of both cell type- and tissue-specific genes would further improve stratification (Figure 4a).

**Figure 4:**
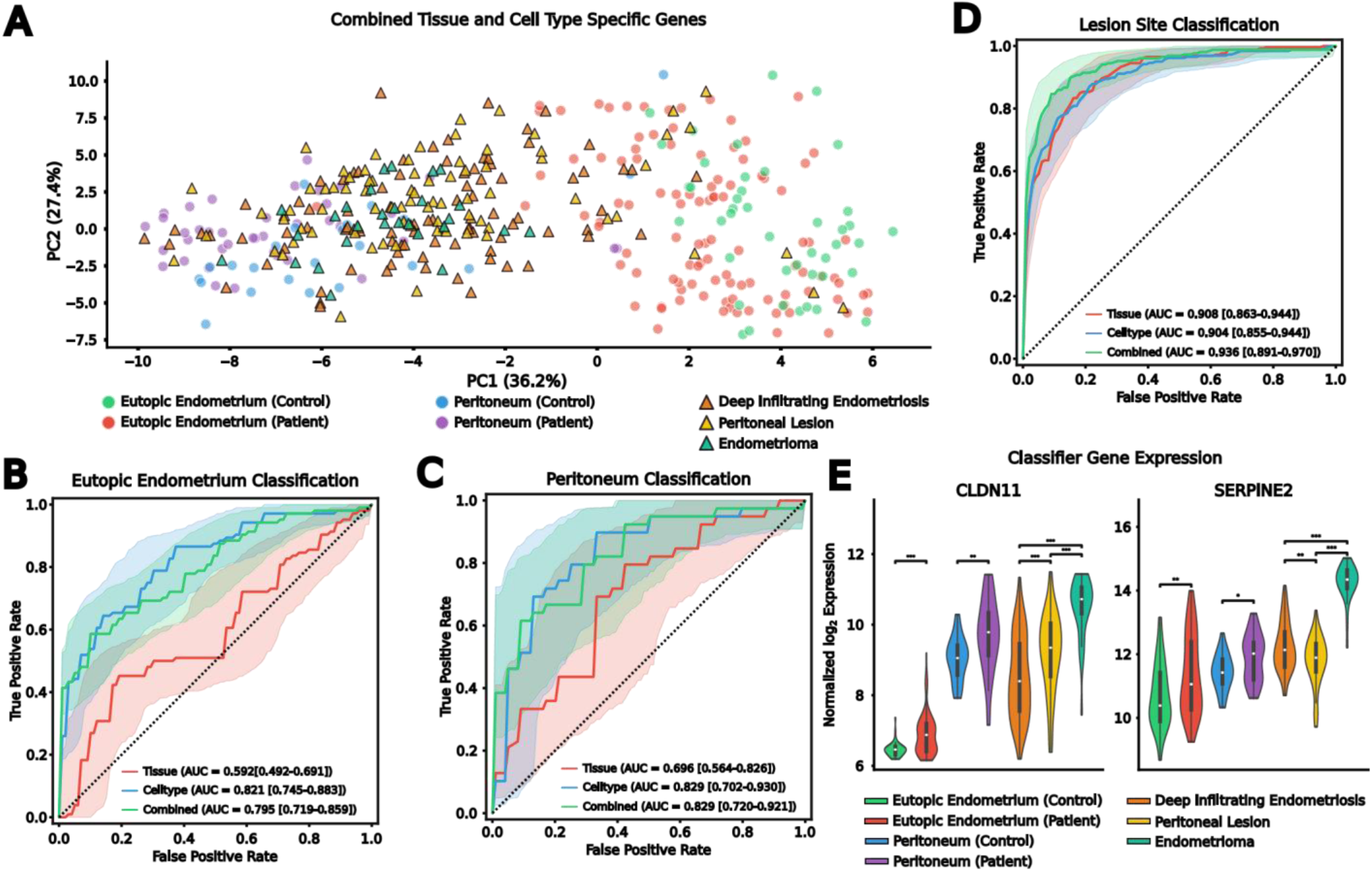
Severity-associated cell type- and tissue-specific signatures classify disease status and lesion origin. **A**, PCA embedding of combined cell type- and tissue-specific gene signature in the bulk tissue micro array data. **B**, Performance of each of the different gene sets to diagnose bulk eutopic endometrium samples as having endometriosis or not as shown via a receiver operator characteristic (ROC) curve. **C,** Performance of each of the different gene sets to diagnose bulk peritoneum samples as having endometriosis or not as shown via a ROC curve. **D**, Performance of each of the different gene sets to stratify lesions by tissue of origin as shown via a ROC curve. **E,** Log normalized expression of *PDLIM3* and *SERPINE2* across all groups. Statistics shown are from a Wilcoxon rank sum test with a Benjamini-Hochberg correction across all possible comparisons for a given gene. * is for a corrected p-value below 0.05, ** is for a corrected p-value below 0.01, and *** is for a corrected p-value below 0.001. All AUC curves show the calculated curve and a 95% confidence interval.

We therefore tested whether these signatures could classify disease status and lesion origin across three clinically relevant comparisons (Methods). We first created a Random Forest classifier which distinguishes between healthy and diseased eutopic endometrium. For this classifier, the tissue-specific genes, which included very few endometrium-specific markers, yielded a statistically insignificant AUC of 0.592 [0.492-0.691], whereas the cell type-specific signature achieved an AUC of 0.821 [0.745-0.883] (Figure 4b), demonstrating that cell type-specific signatures are sufficient to deconvolve endometriosis status in the eutopic endometrium. A similar hierarchy emerged for a classifier built to identify control versus diseased peritoneal tissue, where the tissue-specific classifier achieved a modest AUC of 0.696 [0.564-0.826] and the cell type-specific classifier reached 0.829 [0.702-0.930] (Figure 4c). The improved tissue-specific performance relative to the endometrium comparison suggests a stronger contribution of tissue-level signatures in distinguishing diseased peritoneum from control peritoneum. We then created a third classifier for lesion subtype classification, where we identified if tissue came from a deep infiltrating peritoneal lesion, a superficial peritoneal lesion or an endometrioma, the cell type-specific (AUC = 0.904 [0.855-0.944]) performed worse than the combined tissue and cell type signature (AUC = 0.936 [0.891-0.970]) classifier (Figure 4d), reflecting an increased contribution of tissue-specific genes in this context. Consistent with this, the tissue-specific signature alone was most effective in classifying lesion type of origin (AUC = 0.908 [0.863 −0.944]) (Figure 4d), further reinforcing the presence of an underlying tissue-of-residence signature distinguishing lesion subtypes. Notably, combining cell type- and tissue-specific signatures into a unified gene set yielded minimal gains in classification accuracy compared to just the cell-type specific classifier, consistent with the possibility that cell type-specific genes already encode sufficient tissue-level information to distinguish disease states and lesion subtypes.

Several genes driving these classifiers have established biological relevance in endometriosis. Across both healthy compared to patient tissue- and cell-type-specific classifiers, the stromal cell genes *CLDN11*, previously proposed as an endometriosis biomarker^65^, and *SERPINE2*, a stromal-specific gene, were among the highest-contributing features (Supplementary Figure 9a-c). Both genes showed a stepwise increase in expression from control to patient endometrium and peritoneum, and from these tissues to lesions (Figure 4e), with both genes reaching their highest levels in endometrioma, consistent with the dense stromal compartment characteristic of this lesion subtype. In the eutopic endometrium classifier, the top-contributing genes included immune-specific *CD2*, *GZMH*, and *IL7R* (Supplementary Figure 10a), consistent with prior evidence that an enhanced inflammatory state in the eutopic endometrium may serve as an indicator of endometriosis^66^. The peritoneum classifier also showed high importance of immune cell-specific genes including *RUNX3* and *CCL4L2* (Supplementary Figure 10b), reflecting the established inflammatory environment in the peritoneum of endometriosis patients^67,68^. In the lesion classifier, stromal cell-specific genes such as *SERPINE2* and *CDLN11* (Figure 4e; Supplementary Figure 10c) were among the strongest contributors, consistent with the site-dependent mesenchymal signatures identified in our earlier analyses. Notably, the lesion classifier assigned high importance to *EMX2*, an endometrium-specific gene, with less or negligible importance for some ovary-specific genes (*GREB1* and *OGN* respectively) and the peritoneum-specific gene *CFD* (Supplementary Figure 10c), suggesting that endometrial identity is one of the primary discriminative features for lesion classification.

Altogether, these results demonstrate that cell type- and tissue-specific transcriptomic signatures derived from the integrated single-cell atlas are sufficient to classify diseased from control tissue in both the endometrium and peritoneum, while also capturing the spatial signatures that distinguish lesion subtypes, establishing a molecular framework with potential translational applications for endometriosis diagnosis and subtyping.

## Discussion

Endometriosis affects roughly one in ten women of reproductive age worldwide, yet after nearly a century of research there are still no adequate tools for non-invasive diagnosis^1,2^. This delay is driven in large part by the persistent failure to identify robust non-invasive biomarkers at the blood, urinary, or menstrual effluent level^69^. A fundamental barrier to biomarker discovery is the cellular heterogeneity of ectopic lesions, which comprise an admixture of endometrial-derived cells and host tissue constituents whose signals confound bulk molecular analyses. To address this, we compiled, across multiple data sources, a comprehensive multi-tissue single-cell atlas, across 112 donors, of 672,051 cells across the eutopic endometrium, menstrual effluent, ovaries, peritoneum, and ectopic endometrium from both peritoneal and ovarian sites. This enabled, for the first time, a systematic deconvolution of endometrial-like transcriptomic signatures from the surrounding microenvironment across multiple tissues, hormonal states, and disease severities.

A central finding of this study is that the majority of cells within resected ectopic lesions are transcriptomically more similar to their host tissue of residence than to the eutopic endometrium. This observation carries important implications for how single-cell studies of endometriosis are interpreted. Previous comparisons of transcriptomic profiles between ectopic sites, without first deconvolving the fraction of cells most similar to host tissue, are, in effect, comparisons between two different tissue microenvironments rather than between the most disease-relevant endometrial-like populations. By training cell type-level classifiers on reference tissue signatures and applying them across the ectopic endometrium, we were able to identify the subset of “core” lesion cells which most closely resemble eutopic endometrial cells and to characterize them separately from the host tissue compartment. This analytical framework is conceptually analogous to methods developed for resolving cell-type location in single-cell data from spatially dependent gene expression^70,71^, but applied here entirely within a single-cell context to resolve tissue-of-origin identity.

The identification of core endometrial-like cells at both ectopic sites is broadly consistent with Sampson’s retrograde menstruation hypothesis^22^, which posits that shed endometrial tissue implants at ectopic locations to cause endometriosis. Recent genomic evidence has further supported this model through the identification of shared somatic mutations between eutopic endometrium and matched ectopic lesions^72,73^. Our own analysis of paired eutopic and ectopic samples from the Fonseca et al. dataset similarly identified shared variants between matched tissues, alongside highly concordant mutational spectra dominated by the SBS5 signature indicative of common mutagenic processes. Together, these shared mutations and spectral concordance provide genomic evidence that cells in ectopic lesions descend from uterine precursors, and the observation that variant allele frequencies are elevated in ectopic tissue relative to eutopic endometrium suggests a modest clonal expansion at ectopic sites. The small clonal cell fractions of shared mutations additionally point to an oligoclonal lesion structure, likely reflecting both intrinsic oligoclonality of the seeded population and substantial contributions from the surrounding tissue microenvironment. Our transcriptomic data complement this genomic evidence: the retention of a uterine gene expression identity in core lesion cells, despite their residence in a foreign tissue microenvironment, is precisely what the retrograde menstruation model predicts. At the same time, the fact that the majority of cells at ectopic sites resemble their host tissue rather than the endometrium indicates that lesion establishment involves extensive remodeling and recruitment of local cell populations, consistent with a model in which retrograde-seeded endometrial cells serve as a nidus that co-opts the surrounding tissue environment.

Our classifier-based approach provides a complementary transcriptomic perspective, demonstrating that the endometrial-like fraction within resected tissue is modest, ranging from 5% in endothelial cells to 33% in lymphoid cells, and that the non-core compartment carries distinct transcriptomic programs. Notably, we observed differential immune cell distributions between the core and non-core compartments across ectopic sites: non-core macrophages at ovarian sites were enriched for programs related to cell migration, while those at peritoneal sites showed greater enrichment for proliferative programs. The migration signature in ovarian non-core macrophages is consistent with evidence that endometriosis triggers continuous monocyte recruitment to lesion sites, with macrophages of distinct origins exerting opposing effects on lesion establishment^74^. We also observed differences in cell-cell communication preservation between the eutopic endometrium and the two ectopic sites, likely reflecting the differences in macro-level structural architectures that characterize ovarian endometriomas, which form enclosed pseudocysts through invagination of the ovarian cortex, versus those in peritoneal implants^6,75^. Taken together, these results demonstrate that core endometrial-like cells carry known transcriptomic signatures of endometriosis and provide insight into how the two lesion microenvironments diverge through disease progression.

The eutopic endometrium undergoes dramatic transcriptomic remodeling across the menstrual cycle^12,13,17^, and ectopic lesions have been shown to retain some degree of hormonal responsiveness^6,8,14^. Accordingly, hormonal variation has long been considered a primary driver of transcriptomic heterogeneity in endometriotic tissue, and this assumption has influenced both experimental design and therapeutic development. Our results refine this assumption, demonstrating that the mesenchymal cell types are more transcriptomically similar across hormonal states than across ectopic sites and that hierarchical clustering of differentially expressed genes in an entirely held-out dataset separates EcO and EcP cells from one another regardless of hormonal condition. Critically, however, this site dominance is itself cell-type-dependent and it holds most strongly for the mesenchymal and structural populations, whereas immune cell types retain greater hormonal responsiveness and modulate their functional state with cycle phase. The stability of these site-dependent signatures echoes observations from other biological contexts demonstrating that the local tissue microenvironment is the dominant determinant of cell state, as has been established for specific tissue-resident macrophages^76,77^ and cross-tissue populations of other cell types more broadly^20^. Our findings extend this principle to endometriosis, suggesting that the ectopic tissue niche imprints a durable transcriptomic identity on endometrial-like mesenchymal cells that supersedes hormonal cycling, even as the immune compartment continues to track hormonal state. This compartment split reconciles the dominance of site in subtype discrimination with the hormonally tuned immune programs relevant to cyclical symptom variation.

This hormone-independent site specificity has direct implications for biomarker development. A longstanding challenge in endometriosis diagnostics has been the confounding effect of menstrual cycle phase on candidate biomarker performance, necessitating cycle-matched sampling that limits clinical scalability. Our observation that mesenchymal gene signatures for discriminating between subtypes of endometriosis are conserved across hormonal states suggests that robust biomarkers can, in principle, be identified for endometriosis subtyping without requiring cycle-phase stratification. We demonstrated this by showing that cell type-specific signatures derived from all cell types in combined single-cell atlases are sufficient to classify disease status in independent bulk tissue microarray data for both the eutopic endometrium and the peritoneum, while tissue-specific signatures can only effectively distinguish lesion subtypes. These complementary scales of molecular identity may correspond to distinct diagnostic modalities.

The plasma cell-free transcriptome has been shown to represent a linear combination of tissue-specific RNA contributions^62,78^, with recent work demonstrating that cell-type-of-origin can be resolved from circulating cell-free RNA at single-cell resolution^62^. High tissue-specific classification accuracy for lesion subtype in our analyses suggests that peripheral blood cell-free mRNA could, in principle, be leveraged to identify and classify endometriosis subtypes non-invasively, as has been demonstrated for cancer detection and subtyping^79^. In parallel, endometriosis, by virtue of its transcriptomic signatures in the eutopic endometrium, is uniquely amenable to diagnostic approaches based on menstrual effluent, a non-invasive medium that has shown promise for endometriosis biomarker discovery^18,80,81^. Given that the cell type-specific signatures are sufficient to distinguish diseased from healthy bulk eutopic endometrium, the cell type-specific fraction of cell-free mRNA within menstrual effluent represents a potential avenue for future diagnostic development.

In summary, this study provides a multi-tissue, multi-condition single-cell reference atlas for endometriosis that enables deconvolution of endometrial-like cells from their host tissue microenvironment. We demonstrate that ectopic site, rather than hormonal state, is the dominant driver of transcriptomic heterogeneity among the mesenchymal core lesion cells. The cell type- and tissue-specific signatures identified here are sufficient to stratify disease status and lesion site in independent bulk data, establishing a molecular framework that can be extended to non-invasive diagnostics through peripheral blood cell-free transcriptomics and menstrual effluent analysis. More broadly, this re-analysis paradigm of deconvolving spatial and cell type-specific disease signatures from heterogeneous tissue samples is generalizable to any disease with a resident tissue component, and offers a scalable approach for robust classifier development across heterogeneous patient populations. Given that site-dependent transcriptomic signatures persist regardless of hormonal state, the retrograde menstruation model gains further support from this analysis: lesions at different anatomical sites maintain distinct molecular identities that reflect a combination of their originating tissue and their local tissue context.

## Methods

### Single cell transcriptomics data integration and cell typing

We collected processed scRNA-seq datasets from previously published studies on human endometrial tissue, human ovary tissue, and human endometriosis lesions. These datasets included: the Human Endometrial Cell Atlas (Mareckova et al)^17^, Tabula Sapiens 2.0 Ovary Data^20^, Jones et al^19^, Fonseca et al endometriosis lesion cells and unaffected ovary samples^15^, Tan et al endometriosis lesion cells and peritoneal cells^16^, and menstrual effluent single cell data from Shih et al^18^. The Human Endometrial Cell Atlas also included endometrial biopsy data from the following studies: Wang et al^12^, Garcia-Alonso et al^82^, Tan et al^16^, Lai et al^83^, Fonseca et al^15^, Huang et al^84^. For the endometriosis lesions datasets only studies which included both peritoneal and ovarian endometriosis lesions were included.

We integrated the data from different sources using Harmony^85^ using both dataset and individual as our batch correction variables. The resulting Harmony-corrected principal components were utilized to estimate a neighbor graph and generate a uniform manifold approximation and projection (UMAP) visualization. Following embedding of the samples on a UMAP, we followed the AnnDictionary automated cell type labeling pipeline^21^ using Claude Sonnet 3.5. This gave us preliminary broad cell type labels. These labels, if they contained cell type names which were too broad, such as “T Cell” were further annotated down to CD4, CD8, and T reg identities using the same marker genes used in the HECA atlas. Additional manual cell typing was applied on a lineage-to-lineage basis, where each individual cell type lineage was re-clustered independently and more fine-grained cell typing was performed on the AnnDictionary assigned cell types using literature-based marker gene expression.

### Somatic Mutation Calling and Analysis

Somatic mutations were called de novo from scRNA-seq data from the paired eutopic and ectopic samples found in Fonseca et al 10X 3’ single cell RNA sequencing data using SComatic^86^. In brief, the raw .fastq files, which were downloaded from GEO accession GSE213216, were processed in Cell Ranger 10.0.0, using GRCh38, and produced aligned BAM files. These files were then split according to their cell type annotation and SComatic was used to pileup bases across individual cell types, compare variants to the reference genome, annotate the variants, and filter them. The variant filtering process removed sites which were present in a panel of normals and also removed known RNA editing sites as described previously^86^.

To assess selection pressure on coding mutations we computed the ratio of nonsynonymous to synonymous mutations (dN/dS) using the amino acid consequences of the observed mutations, by intersecting variant positions with GENCODE CDS annotations from GRCh38. To generate a null expectation under neutrality, we performed a permutation test across 100,000 iterations. In each permutation, at each of the 213 coding variant positions, a random alternative base was sampled with probabilities weighted by the observed trinucleotide spectrum at the mutation’s context. This sampling was performed across all mutation sites followed by the calculation of the dN/dS. Significance was assessed by computing a z-score and a p-value from the standard normal distribution.

To identify shared mutations between eutopic and ectopic samples we used the pooled variants at each patient-tissue pair to identify the same somatic changes. To further identify the clonal cell fraction, the percentage of cells carrying a given mutation, we ran the de novo somatic variant calling on the cell type BAM files independently, without bulking across cells. This resulted in high confidence mutation calling for a select group of cells, these high confidence single cell mutation calls, were then used to estimate the fractional contribution of the shared mutations to the overall population of cells. The pooled variants across our samples were utilized to call both the variant allele frequency and accumulation of mutations per cell type.

### Single cell transcriptomics-based endometrium classifier

After integrating the full single cell object we trained cell type specific binary classifiers to identify the most endometrium-like cells in the lesions. To train this model we utilized the reference data from the HECA, menstrual effluent, ovarian datasets, and peritoneum data from the Tan et al study. For each annotated cell type we separated our data into the cells from the eutopic endometrium, or menstrual effluent, and those from another tissue source. Cell types were included only when both classes had 200 or greater cells per class.

We created ElasticNet classifier models trained on log-normalized gene expression for each cell. Features were selected in a fold-specific manner to avoid information leakage. Within each training split, we performed differential gene expression analyses with a Mann-Whitney-Wilcoxon test comparing the endometrium to all other tissues. We filtered the genes for the classifier by Benjamini Hochberg adjusted p-value, minimum absolute log2 fold-change (1), and minimum detection frequency (20%). To reduce the influence of broadly expressed or ambiguously annotated features, we excluded mitochondrial and ribosomal genes and additional low-specificity gene families and identifiers using a fixed pattern-based filter. We also included the opportunity for a tissue-specificity filtering step for the differentially expressed genes, via a cell type level tau calculation^87^ across the different tissues. This was computed from tissue-wise mean expression within the training split.

Classifiers were ElasticNet models which tuned regularization strength and sparsity by grid search over multiple values of the inverse regularization parameter and the ElasticNet L1 ratio. Performance was evaluated using nested cross-validation with donor-disjoint outer folds to ensure that training and testing sets contained non-overlapping donors. Hyperparameters were selected within each outer training fold using an inner cross-validation loop, and the inner folds were also donor-aware. To mitigate donor imbalance in some configurations, we applied donor-balanced sample weighting within training folds, reweighting cells inversely proportional to the number of cells contributed by each donor within each class.

To make the downstream calls equivalent to a probability, we calibrated predicted probabilities using sigmoid calibration fit on the training data within each outer fold, and then generated calibrated probabilities for the held-out validation split. For each outer validation fold, we quantified discrimination using area under the precision–recall curve and area under the ROC curve. We additionally reported balanced accuracy, F1 score, and confusion matrices using an 80% probability cut off. For each configuration and cell type, we trained a final model on all available cells using the same feature-construction strategy, tuned hyperparameters using donor-aware cross-validation when possible, and serialized the fitted model together with the selected feature list and cross-validation summary.

Selection of the final model was performed through a parameter sweep across feature strategies (differential-expression genes with or without tau filtering), feature-set sizes, filtering thresholds, and negative-class definitions. To select a single model per cell type, we aggregated cross-validation metrics across runs and ranked candidates. We prioritized configurations meeting a minimum F1-performance threshold of 0.9 in the tau constrained models, using non-tau filtered models as a fallback when the tau-based models did not meet the minimal F1 threshold. Selected runs were consolidated into a final model set and summarized with diagnostic plots comparing performance, calibration, and feature-set size across cell types.

### Differential Gene Expression

Differential expression was performed using a two-part hurdle model (MAST)^88^ to identify transcriptional changes within each cell type. For each cell type lesion cells were stratified using the classifier derived probability, lesions were divided into cells above or below a fixed probability threshold of 80%. For comparisons involving eutopic endometrium and lesion cells, reference endometrium was additionally partitioned to just the samples having endometriosis. For severity-stratified analyses, samples were categorized into mild or severe disease based on stage groupings, and comparisons were performed within severity strata.

To minimize confounding by menstrual cycle state across donors, differential expression was performed using menstrual cycle stage as a covariate. Cells were grouped into discrete cycle-state bins, and only cycle states represented in both case and control groups were analyzed. Differential expression was then run independently within each cycle-state subset. For each gene, we extracted a single, combined significance value from the hurdle component of the model to test for differential expression between case and control. Effect sizes were reported as log fold changes from the combined model and converted to log2 scale. Multiple testing correction was performed using the Benjamini-Hochberg procedure to control the false discovery rate. For each run, the output table included per-gene log2 fold change with confidence intervals, test statistics and P values, and FDR-adjusted P values.

### Comparing core and non-core cells to the eutopic endometrium

To first compare the core and non-core cells of the lesion cells in different sites to the eutopic endometrium, we performed a stratified analysis by performing differential communication and differential expression analysis. The differential communication analysis was performed with scDiffComm^28^, in which both the ectopic peritoneal and ovarian core and non-core cells were compared to the eutopic endometrium for a total of 4 separate comparisons. Then using the output tables from scDiffComm we isolated the “FLAT” interactions, those which were not changing from the eutopic endometrium, with an FDR adjusted p-value of below 0.05 and an odds ratio (OR) of greater than 1. Then to compare the core versus non-core interaction changes within each of our lesion types we subtracted the ORA Score, combined adjusted p-value and log2(OR), of the non-core cells from the core cells to identify stronger FLAT interactions in the core cell or non-core cell populations.

Additionally, we compared the core and non-core cells to the same cell types in the eutopic endometrium of individuals with endometriosis using the strategy outlined above. We were then able to make cross core and non-core comparisons by looking for unique genes. We also took the Jaccard index between these core-specific gene sets for every cell type across the two separate ectopic sites. Finally, utilizing gseapy, we searched for the biological process gene ontology terms in the differentially expressed genes for specific lineages across the sites. This was performed, in brief, by taking the union of the different lineage differentially expressed genes, running the gseapy enrich function, and identifying gene programs in which 3 or more genes were present and removing gene sets that did not contain at least a 10% overlap of gene terms, and removing gene sets which were redundant with 50% or more of overlapping genes.

### Identifying Hormone Sensitive Gene Modules

In order to identify gene modules in endometrial cells which are responsive to different stages of the menstrual cycle and hormone therapy we divided our dataset into tissue-cell type pairs, where tissue is either eutopic, the core endometrial peritoneal lesion cells or the core endometrial ovarian lesion cells. We then tested for differentially expressed genes across hormonal states (Proliferative, Secretory, or Exogenous Hormones) using MAST, with donor included as a fixed effect. Cycle responsive genes were identified by thresholding the genes for a Benjamini-Hochberg adjusted p-value of less than 0.05 and a maximum log2 fold change of greater than or equal to one, where the peak-versus-trough fold-change was computed as the difference between the highest- and lowest-expressing phases for each gene.

To identify recurrent trajectory programs across the hormonal status which span the eutopic and ectopic tissue we pooled all gene-tissue-cell type triplets. The mean expression of the cycle-responsive genes for these triplets was then z-scored and subject to k-means clustering. We swept the number of clusters from 3 to 10, where k=6 yielded the highest silhouette score (0.540). To annotate each cluster with its dominant biological program, we took the union of genes whose triples were assigned to that cluster and ran gene set enrichment analysis for biological processes using gseapy. The resulting term list was filtered to adjusted p-value < 0.05 and greater than or equal to 2 overlapping genes, then a greedy redundancy filter was applied that iteratively selected terms by adjusted p-value while excluding terms whose gene content overlapped previously-selected terms by more than 50%. The top three non-redundant terms per cluster were used as cluster labels.

For each cell type, cluster-assigned genes were partitioned into mutually-exclusive site categories based on which tissue contexts they were called cycle-responsive in: eutopic-only, EcO-only, EcP-only, or shared (assigned in two or more tissues). Per-celltype site-partition counts were visualized as stacked bars. We then identified the clusters which reached a peak or had their minimum in the Secretory phase of the menstrual cycle. For each cluster group, we identified genes whose triples were assigned to that cluster in at least one ectopic tissue (EcO or EcP) but not in eutopic, the ectopic-only gene set per cell type. Ectopic-only genes from EcO and EcP were pooled per cell type, and gene set enrichment analysis was run separately for the secretory-peak (C3) and secretory-trough (C1 or C6) pooled gene sets. Cell types with fewer than two genes in the ectopic-only set were excluded from enrichment.

### Core Endometrial Cell Modules

To identify co-expressed gene modules across the core endometrial cells of both ectopic sites, consensus non-negative matrix factorization (cNMF) was applied to the raw gene expression count matrix across matrix ranks ranging from 10 to 55, with 1000 iterations per K. The optimal number of factors, K = 33, was determined by evaluating the trade-off between reconstruction error and stability across K values. All factorization outputs from the 1000 iterations at K = 33 were aggregated and subjected to density filtering, yielding 33 robust gene modules.

### cNMF Analysis

To interpret cNMF programs, we performed pathway enrichment on the top program genes using gseapy. For each cNMF factor, the top 100 genes with the highest weights were selected for gene ontology (GO) analysis. For 100 genes in each cNMF factor, pathways with an adjusted p-value < 0.01 were retained. Among these, up to 10 pathways were selected per factor, a candidate term was discarded if more than 25% of its overlapping genes were shared with any single higher-ranked term retained for the same factor. We further enforced global non-redundancy by discarding terms whose overlapping gene sets shared more than 25% of their genes with a term already retained for a different factor. Because these filters operate on leading-edge gene sets rather than on term identity, the same GO term may be retained for more than one factor when the underlying genes are distinct. This procedure returned 47 terms across 26 of the 33 factors, with one to four terms per factor. Where a factor was annotated with multiple terms, the term with the lowest adjusted P value was used as its representative label. The complete mapping is provided in Supplementary Data Table 1. Annotated terms and associated gene sets were compiled into a program-to-pathway mapping used for labeling downstream visualizations and summarization.

To summarize cell-type specificity, each cell’s program usage vector was normalized to sum to one across all 33 programs. We then computed mean normalized usage of each annotated program within each of 19 cell types and z-scored these means across cell types within each program (Figure 3c). Each program was assigned to the cell type at which its z-scored usage was maximal. For downstream analysis we retained only programs whose usage was specific to a single cell type, defined as a margin of at least 0.5 standard deviations between the highest and second-highest cell-type z-score; 22 of the 26 annotated programs met this criterion. Each cell type was allocated a number of programs equal to the number of specific programs assigned to it. Three cell types had no specific program and were instead allocated the annotated program with the highest raw mean usage within that cell type; for two of these, the resulting program was already assigned to another cell type and is shown under both. This yielded 25 cell-type-program pairs spanning 23 unique programs (Supplementary Data Table 2). Because specificity is defined by relative enrichment across cell types whereas the fallback rule uses absolute usage within a cell type, one program is placed with stromal cells in Figure 3c and with glandular epithelial cells in Figure 3d.

To compare program activity across biological contexts, we defined condition labels as the combination of lesion site class (EcO or EcP) and cycle-state bin. Within each cell type and condition, we computed mean normalized usage of each program across the cells of the corresponding cell type within each of the six conditions and plotted as raw mean usage. We tested for differences in program usage between lesion sites while adjusting for hormonal state and accounting for within-donor correlation. For each cell-type-program pair, we fit an ordinary least squares model predicting per-cell program usage from lesion site and hormonal state (usage ∼ site + state) on the cells of that cell type, using cluster-robust standard errors with donor as the clustering variable. Because the model is additive, the site coefficient estimates a single state-adjusted site effect; the reported contrast is EcO minus EcP, with positive values indicating higher usage in EcO. Note that clustering adjusts standard errors for within-donor correlation but does not remove donor-level confounding of the point estimates, as donor is nested within lesion site. Multiple-hypothesis correction was performed using the Benjamini-Hochberg procedure across all real P-values and results are contained within Supplementary Data Table 3.

To quantify global transcriptional differences between conditions while accounting for cell-type composition, we computed pairwise distances in program-usage space using a cell-type–aware centroid approach. For each cell type and condition, we computed the centroid of selected program usage vectors (mean across cells) and calculated pairwise condition distances within each cell type using mean absolute difference across program dimensions. We then aggregated distances across cell types using the mean values across the conditions. To summarize site differences within each cycle state, we extracted the cell type specific distances between matched site pairs within each state and compared distances across states using paired Wilcoxon signed-rank tests across cell types, followed by Benjamini Hochberg correction.

### Hierarchical clustering of hormonally controlled DEGs

To assess the ability of differentially expressed genes in one hormonal state to stratify lesion subtype in other states we first removed the Fonseca et al dataset from the overall single cell object. We then performed differential expression analysis on the core lesion cells in the peritoneal endometriosis samples against the core lesion cells in the endometrioma samples in the Tan et al hormonally controlled samples. We then restrict the differentially expressed genes using an adjusted p-value threshold of 0.01 and a log2fc threshold of 0.75. Amongst these genes for each of our mesenchymal cell types, we isolated the top 15 positive and negative direction genes for each of the 4 cell types, including Stromal Cells, Vascular Smooth Muscle Cells, Fibroblasts, and Myofibroblast like cells. We then observed the expression of these genes on the scaled log normalized expression data from the Fonseca et al core lesion cells across a multitude of menstrual cycle stages. We then applied hierarchical clustering to the pseudobulk expression for donor-tissue-celltype groups and visualized the dendrogram and the heatmap together.

### Creation of tissue and cell type specific gene sets

To compare differential expression patterns across both disease severity and tissue groupings we calculated differential gene expression between reference ovarian tissue and our non-core endometrioma cells, then between reference peritoneum tissue and our non-core peritoneal lesion cells, and between the eutopic endometrium of patients with endometriosis and the core endometrioma or peritoneal lesion cells. If a gene for a comparison had an adjusted P value of less than 0.05 it was kept for effect size analysis. To identify endometrioma lineage based effect sizes, we grouped the individual cell types in both endometrioma based comparisons by lineage and calculated the 250 largest magnitude differential expression values. This process was repeated for both severity classes and the same was repeated for the peritoneal based differential expression values. The lineage level comparisons were performed using a Mann-Whitney-Wilcoxon test with a BH correction of the P values.

To assess whether severity strata shared similar transcriptional responses, we filtered the differentially expressed genes to only include those with a log2 fold change of magnitude 1 or greater and computed the intersection between the mild and severe differential expression genes for each cell type in each comparison in a direction aware manner. We then took the union of the DEG sets across all cell types within each severity grouping and plotted one venn diagram for each tissue comparison to show a broad mild to severe gene overlap.

To interpret the ability of the intersecting genes to be deconvolved in a non-invasive manner we quantified the number of tissue enriched/enhanced genes (HPA)^26^ which also had a Gini coefficient of >0.6, and cell type specific gene sets as defined by the Gini coefficient of >0.8 for differentially expressed genes between cell types in Tabula Sapiens v.2.0 as described previously^62^. For tissue specificity, we only included genes which were specific for the endometrium or ovaries, and the peritoneum specific set was approximated as being tissue specific genes for tissues in the peritoneal cavity, such as the bladder, fallopian tubes, adipose tissue, and intestines. For each intersection, we recorded whether the gene was cell type specific for any of its mapped cell types. To directly compare tissue and cell type specificity within the shared mild to severe genes we classified each gene as tissue or cell type specific. For each comparison, we visualized category proportions using pie charts. We then saved the union of the cell type specific gene across all of our comparisons, we also saved the union of tissue specific genes across all comparisons for downstream analysis.

### Machine Learning Classification of Endometriosis Lesions

Micro-array data was obtained from the Gene Expression Omnibus (GEO accession: GSE141549)^59^, comprising batch-corrected, normalized microarray expression profiles from 205 samples across seven tissue categories: control endometrium, patient endometrium, control peritoneum, patient peritoneum, deep infiltrating endometriosis, peritoneal lesions, and ovarian endometrioma. We employed a patient-aware data partitioning approach during model evaluation, making sure no individual donor was included in both the training and the validation samples.

We created three separate models for every comparison, a tissue-enriched gene set model (described above), a cell type specific gene set model (described above), and a combined gene set model. Genes not present in the expression dataset were excluded, and duplicate gene identifiers were resolved by retaining the first occurrence. We first calculated the first and second principal components of the micro-array data for the three separate gene sets and observed visualization across the different tissues present.

We then created machine learning models for each of our gene sets, using a Random Forest classifier, to distinguish between (1) control and patient endometrium, (2) control and patient peritoneum, and (3) between deep infiltrating endometriosis, peritoneal lesions, and ovarian endometrioma. To train the model features were standardized to zero mean and unit variance prior to model fitting. Hyperparameters were tuned via grid search and the class weights were also balanced to account for unequal class frequencies. Nested cross validation was performed using stratified group k-fold cross-validation (k=5) on both the inner and outer folds, ensuring all samples from a given patient remained within the same fold. Hyperparameter selection was performed via nested cross-validation where the inner loop was used for identifying optimal parameters and the outer loop evaluation providing performance estimates.

Model performance was assessed using the area under the receiver operating characteristic curve (ROC-AUC) for the out-of-fold prediction generated from the nested cross-validation. For our multi-class classifier a macro-averaged one-versus-rest ROC curve was computed for every gene set. Confidence intervals for ROC curves and AUC values were estimated via block bootstrapping with the patient as the resampling unit. In short, for each of the 1,000 bootstrap iterations, *N* patients were drawn with replacement from the *N* unique patients in the analysis, and all out-of-fold predictions belonging to the sampled patients were aggregated to form a bootstrap sample (so a patient drawn twice contributed all of their samples twice). The 2.5th and 97.5th percentiles of the bootstrap distribution defined the confidence bounds. The coefficients of a Random Forest model trained across the entire dataset were then used to rank feature importance.

## Data Availability

All data utilized in this experiment was from publicly available repositories which did not require approval. The HECA was downloaded from a web portal which originated from its parent publication: https://www.reproductivecellatlas.org/endometrium_reference.html. Shih et al menstrual effluent data was available through National Center for Biotechnology Information/Gene Expression Omnibus (GEO) accession number GSE203191. Tan et al lesion and peritoneal data were available through a web portal: https://singlecell.jax.org/datasets/endometriosis-2022. Fonseca et al lesion and ovary data were available through GEO accession number GSE213216. Jones et al single cell ovary data was available through GEO accession number GSE260685. Tabula Sapiens 2.0 could be accessed through the CellxGene Web portal: https://cellxgene.cziscience.com/collections/e5f58829-1a66-40b5-a624-9046778e74f5.

## Code Availability

All code for the work and the manuscript are available on GitHub at: https://github.com/doughenze/endometriosis_sites.

## Acknowledgments

The authors thank all members of the Quake Lab for valuable feedback and discussions throughout the study.

## Author Information

**Department of Bioengineering, Stanford University, Stanford, CA, USA**

Douglas E. Henze, George Crowley, and Stephen R. Quake

**Department of Applied Physics, Stanford University, Stanford, CA, USA**

Stephen R. Quake

## Author’s Contribution

D.E.H, G.C., and S.R.Q conceptualized and designed experiments. D.E.H collected data. D.E.H and G.C analyzed the data. D.E.H and S.R.Q discussed and interpreted the data. D.E.H designed the figures. D.E.H and S.R.Q wrote the manuscript. D.E.H and S.R.Q supervised, directed, and managed the study. All authors discussed the results and commented on the manuscript.

## Competing Interest

The authors declare no competing interests.

## Correspondence

Correspondence to Stephen R. Quake

## Supplementary Figures

**Supplementary Figure 1:**
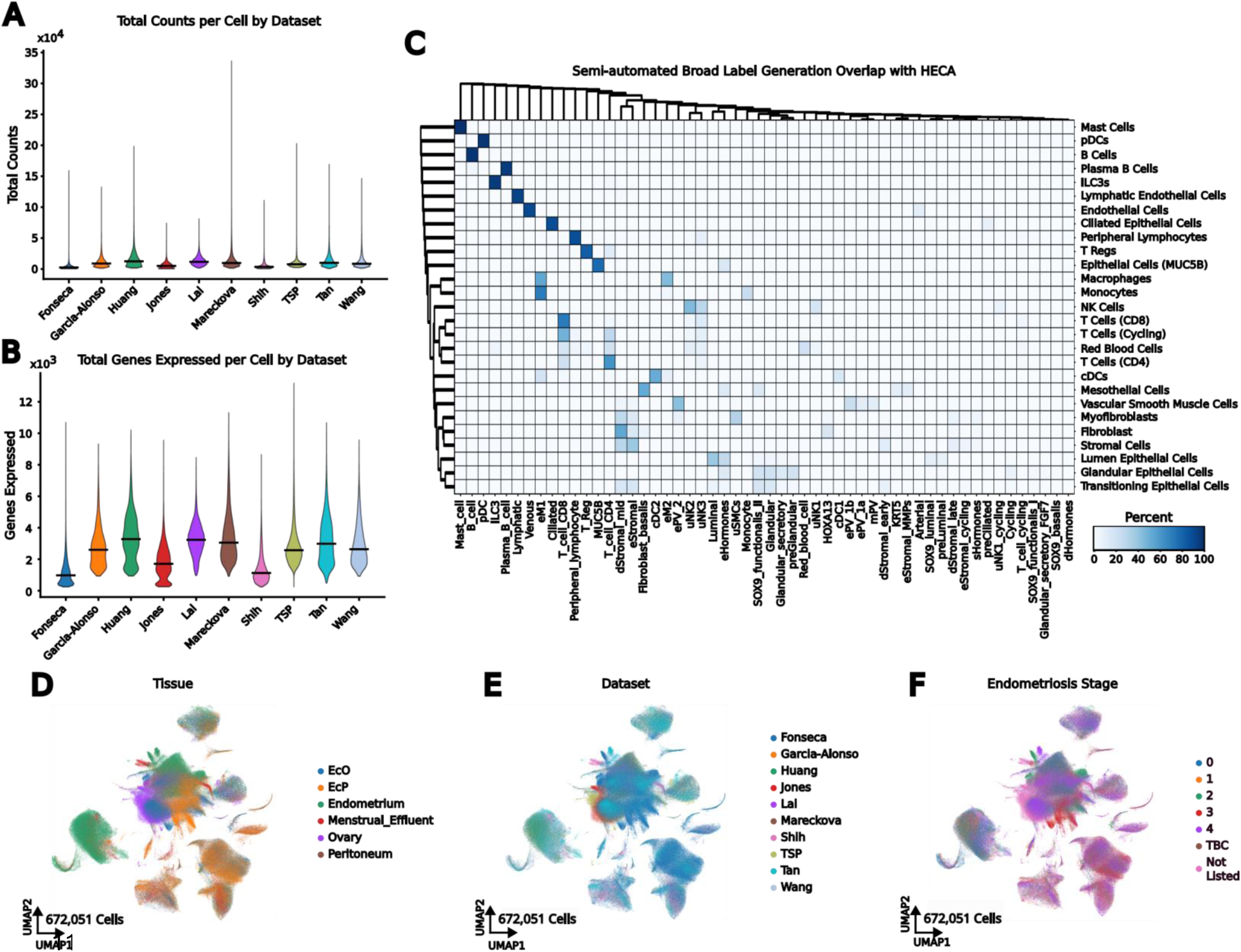
Semi-automated cell typing recovers previously identified cell populations. **A,** Number of transcripts per cell, line on the violin plot indicates the median value. **B,** Number of genes per cell, line on the violin plot indicates the median value. **C,** Cluster map showing the relative percentage of HECA labels for each of our semi-automated pipeline created labels. **D,** UMAP representation of all cells colored by tissue. **E,** UMAP representation of all cells colored by Dataset. **F,** UMAP representation of all cells colored by endometriosis stage.

**Supplementary Figure 2:**
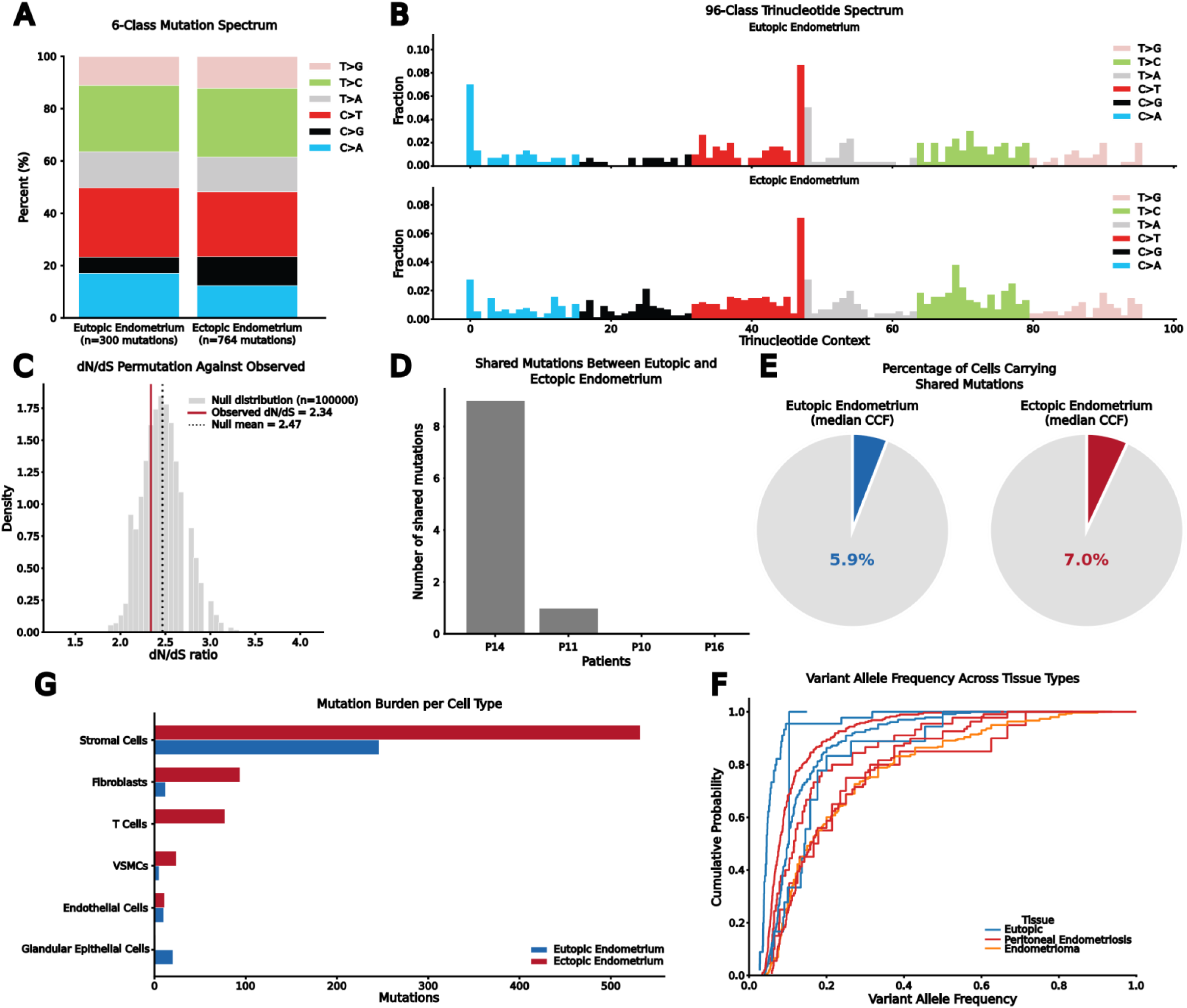
Somatic mutation accumulation in eutopic and ectopic endometrium. **A-B**, 6-class (**A**) and 96 class (**B**) mutation spectra between the eutopic endometrium and the ectopic endometrium. **C**, Permutation test results showing expected against null dN/dS for the observed mutation spectra and mutation sites. **D**, Number of mutations shared between the eutopic and ectopic endometrium in a given patient. **E**, Percentage of cells, bulked across the two patients with shared mutations, that display the shared mutation in the eutopic (left) and ectopic (right) endometrium. **F**, cumulative density functions of the variant allele frequencies across the different patients eutopic and ectopic samples. **G**, Total number of mutations attributable to each cell type, colored by sample location.

**Supplementary Figure 3:**
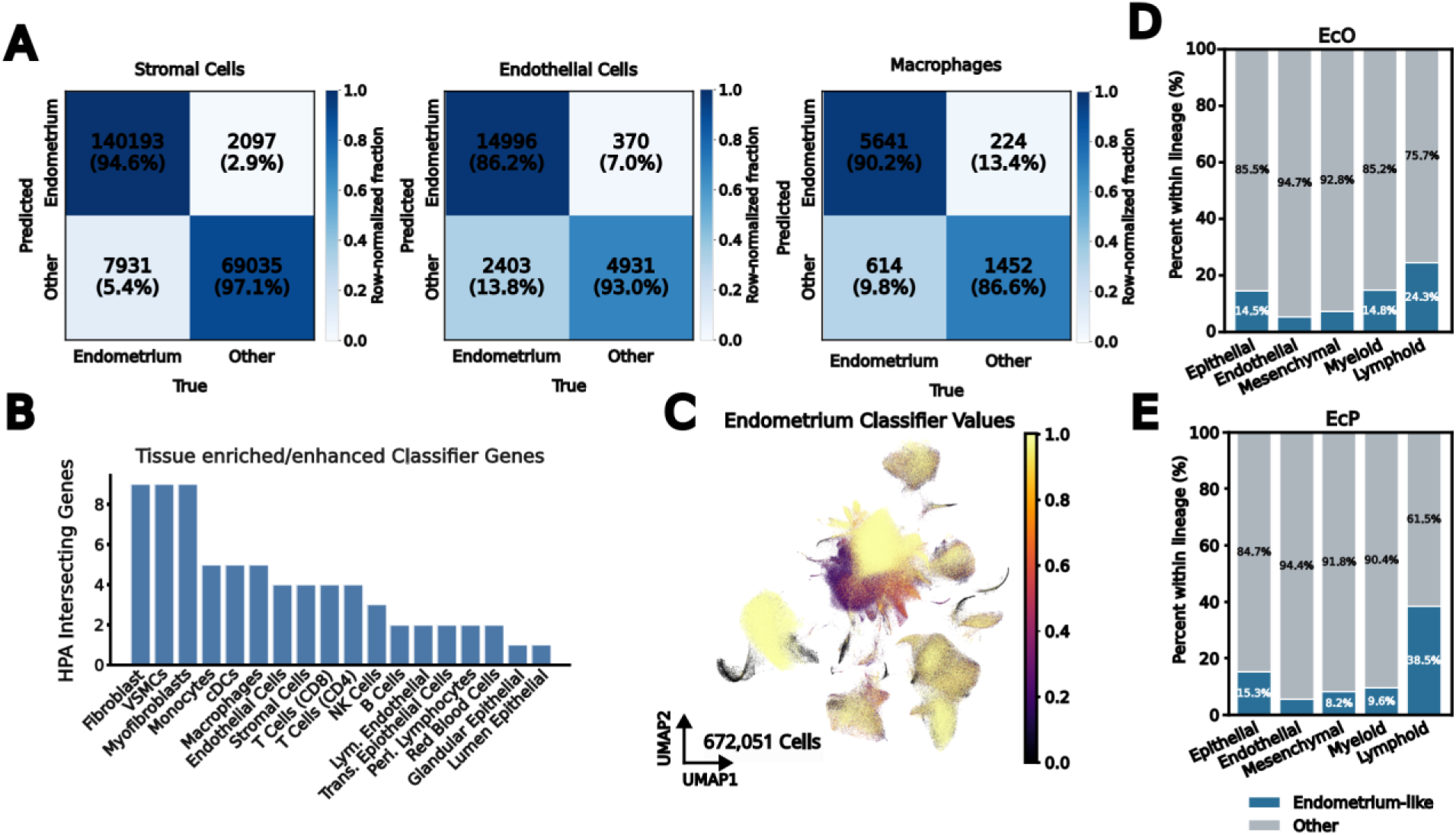
Validation of cell type-level endometrial origin classifiers. **A,** Confusion matrices of classifier predictions across populous cell types **B,** Bar plot showing the number of endometrium tissue-enhanced or tissue enriched HPA genes which were used in the classifier for each cell type. **C,** UMAP showing value of applied classifier across the whole integrated dataset. **D-E,** Stacked bar plots showing classifier values across different cell lineages in the **(D)** endometrioma and the **(E)** peritoneal endometriosis samples.

**Supplementary Figure 4:**
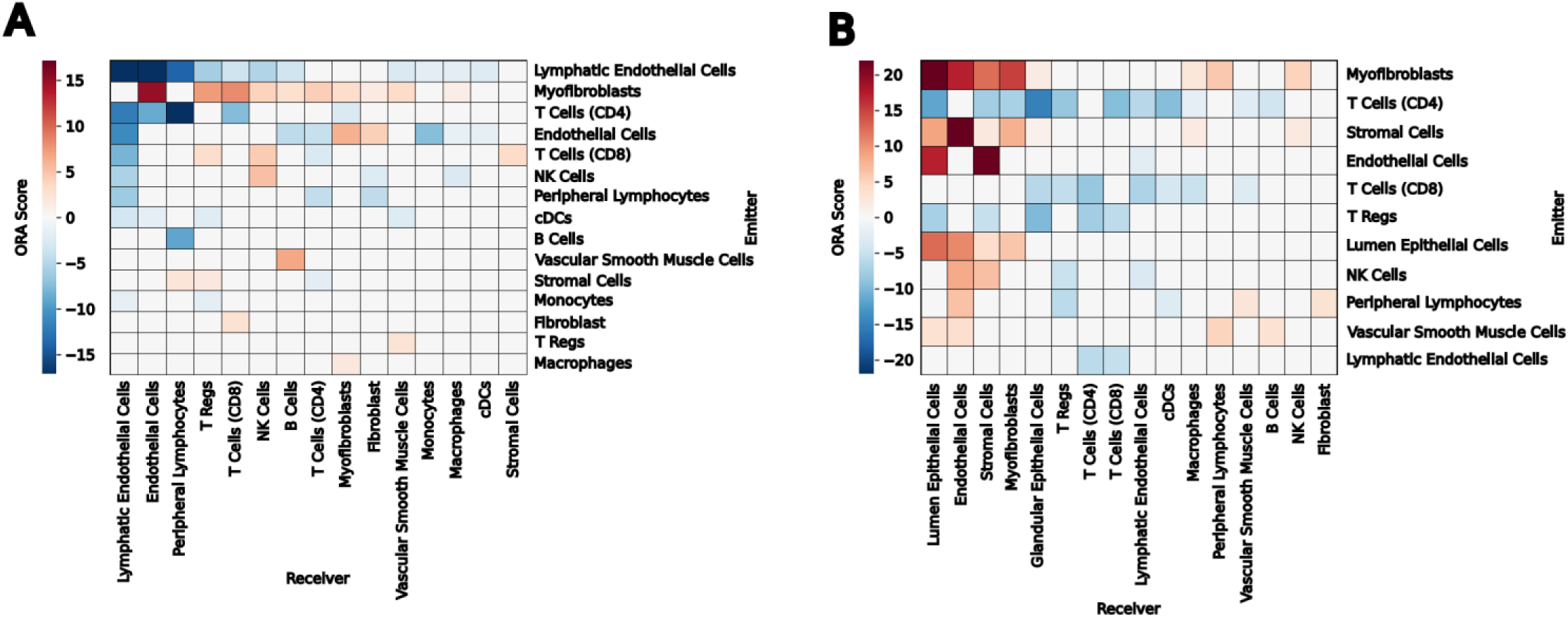
Differential cell-cell interaction between the eutopic endometrium and core or non-core cell populations. **A,** Heatmap showing the difference in odds ratio scores of preserved cell-cell interactions from the eutopic endometrium to core and non-core cells in endometrioma. **B,** Heatmap showing the difference in odds ratio scores of preserved cell-cell interactions from the eutopic endometrium to core and non-core cells in peritoneal endometriosis.

**Supplementary Figure 5:**
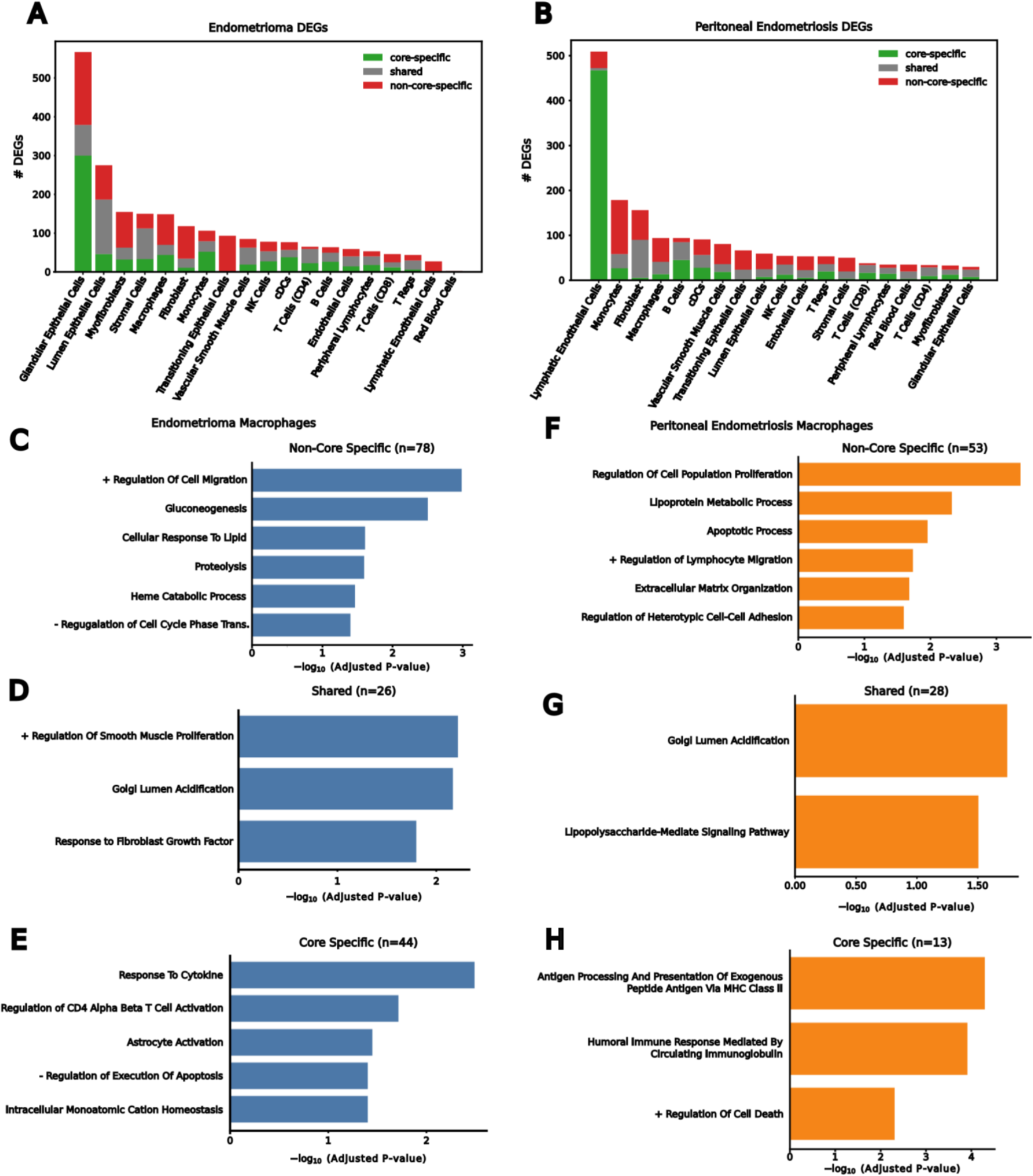
Core- and non-core-specific differential gene expression between the eutopic endometrium and different lesion sites. **A,** Summary of differentially expressed genes as core-specific, non-core-specific, or shared for endometrioma samples across all cell types. **B,** Summary of differentially expressed genes as core-specific, non-core-specific, or shared across peritoneal endometriosis samples across all cell types. **C-E**, Gene Ontology Biological Processes for endometrioma macrophage (**C)** non-core specific, (**D)** shared, and (**E)** core-specific differentially expressed genes. **F-H** Gene Ontology Biological Processes for peritoneal endometriosis macrophage **(F)** non-core specific, **(G)** shared, and **(H)** core-specific differentially expressed genes.

**Supplementary Figure 6:**
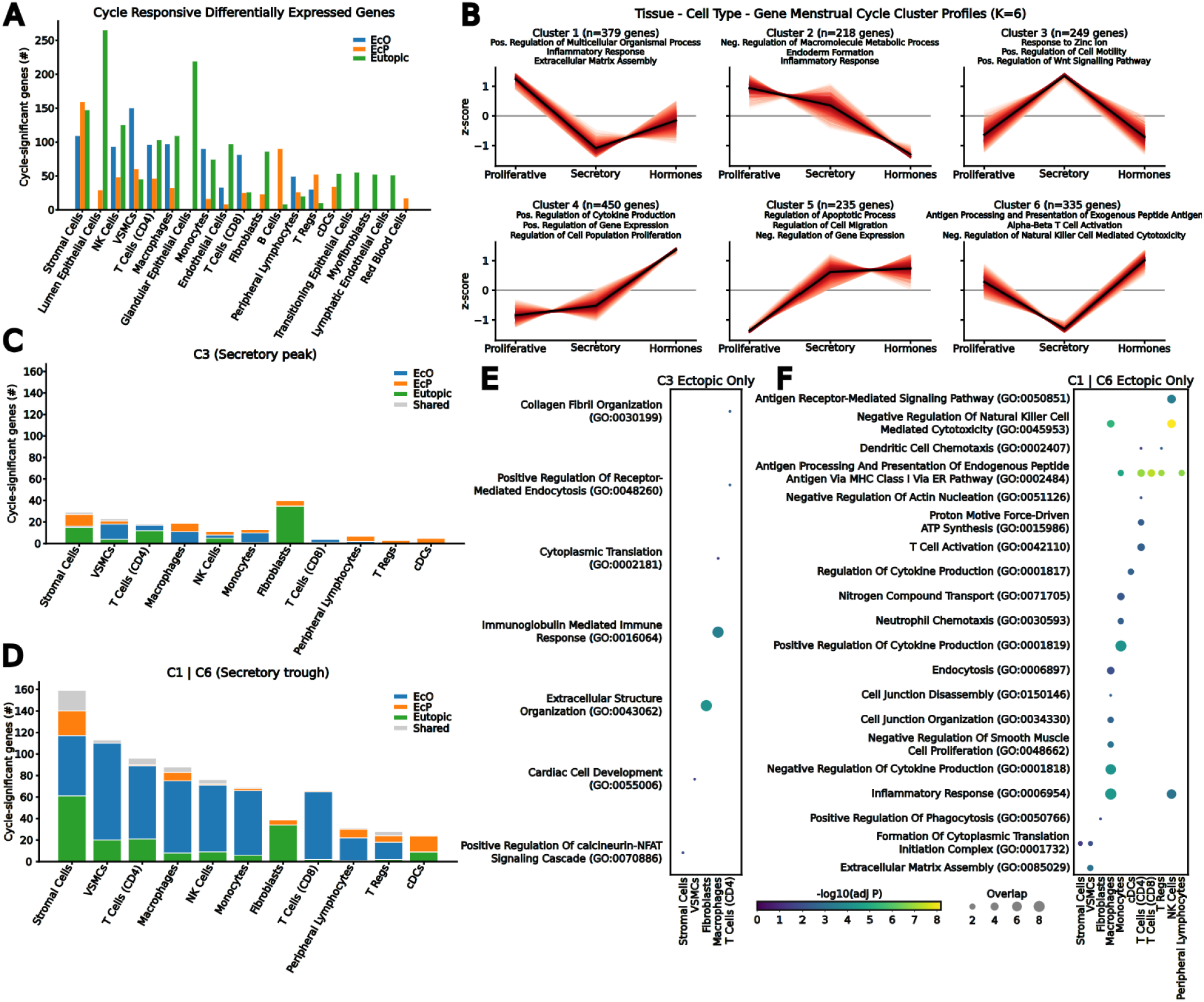
Hormone-responsive gene modules explain endometriosis pain. **A,** Barplot showing the number of cycle-dependent differentially expressed genes for each cell type across the ectopic ovary (EcO), ectopic peritoneum (EcP), and eutopic endometrium (Eutopic) tissues. **B,** Hormone responsive cluster profiles identified through k-means clustering of the mean expression for a tissue, cell type, and gene triplet at each hormonal state. Clusters are labeled with the top 3 gene ontology terms for the union of all genes present in the cluster. **C,** Stacked bar plot representing the number of cycle-responsive genes in Cluster 3 (C3) for EcO, EcP, Eutopic, and shared between two or more. **D,** Stacked bar plot representing the number of cycle-responsive genes in Cluster 1 or Cluster 6 (C1 | C6) for EcO, EcP, Eutopic, and shared between two or more. **E,** Bubble plot showing gene ontology biological processes enriched in the ectopic only (EcO or EcP) genes for Cluster 3 across each cell type. **F,** Bubble plot showing gene ontology biological processes enriched in the ectopic only (EcO or EcP) genes for Cluster 1 or Cluster 6 across each cell type.

**Supplementary Figure 7:**
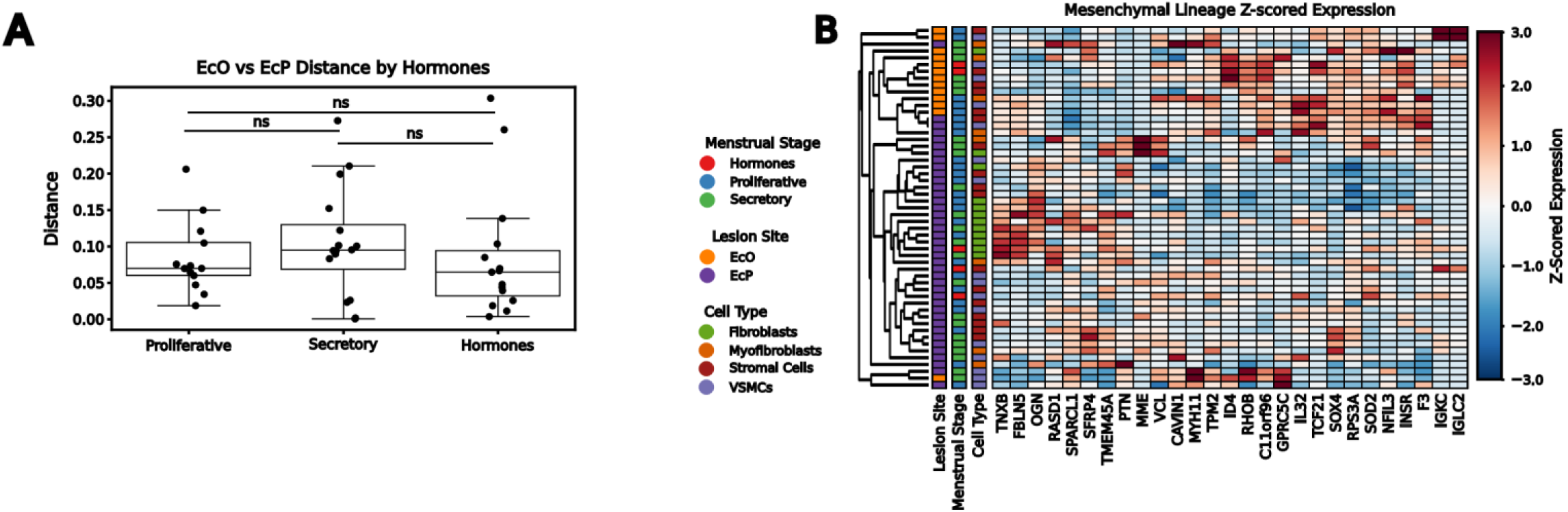
Site-dependent cell type signatures are conserved across the full spectrum of hormonal conditions. **A,** Boxplot showing the euclidean distance between the usage space for each individual cell type across sites for each hormone state. n.s represents no statistically significant deviation. **B,** Clustered heatmap showing z-scored expression of top 15 differentially expressed genes across mesenchymal cell types in a hormonally controlled dataset as applied to a non-hormonally controlled dataset.

**Supplementary Figure 8:**
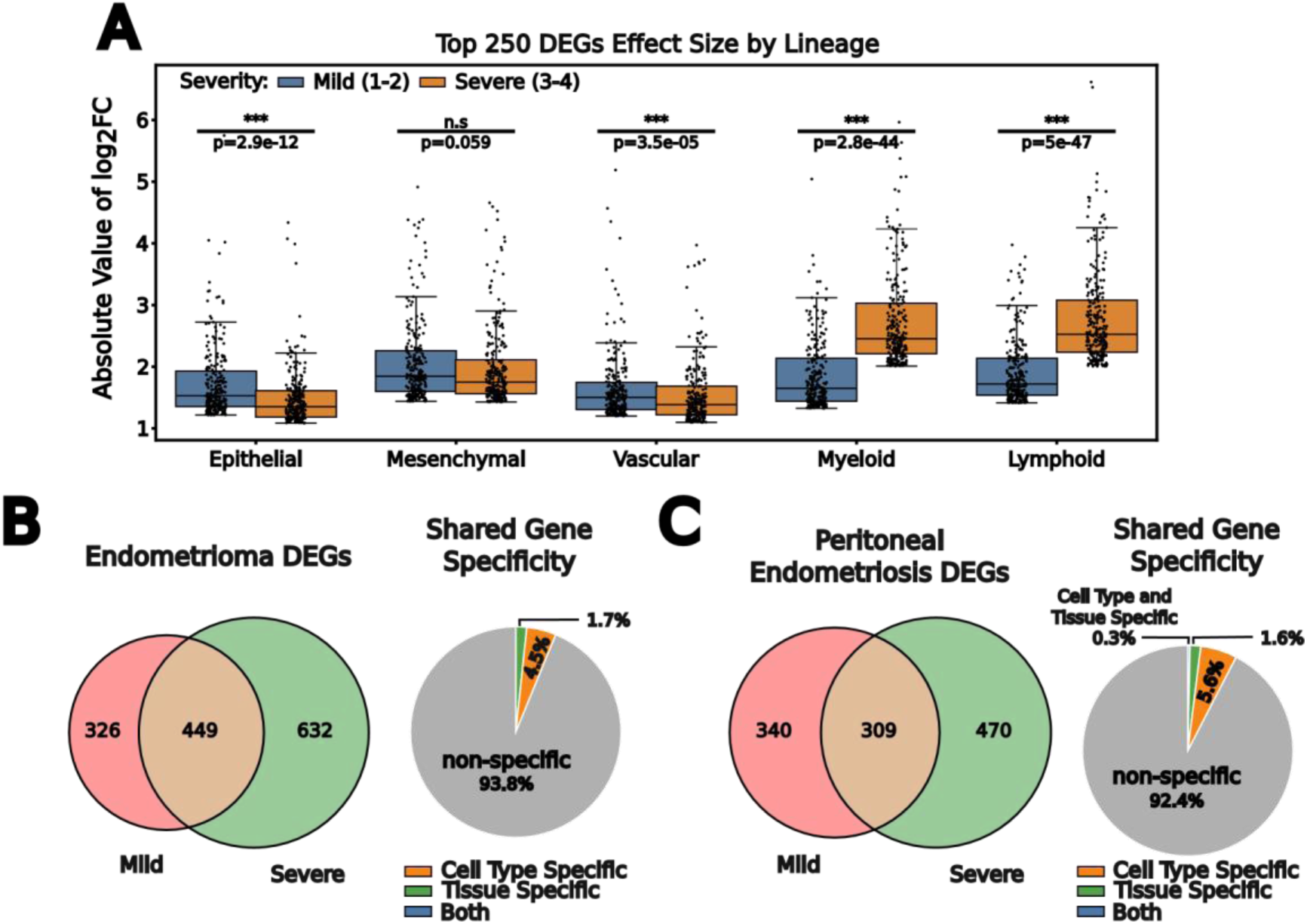
Severity associated differentially expressed genes contain cell type- and tissue-specific genes. **A,** Boxplot representation of the top 250 genes associated with each lineage across different severity classifications of endometriosis. **B**, Venn diagram representation of the differentially expressed genes split by severity across endometrioma (left) and percentage of severity shared genes that are non-specific, cell type-specific, tissue specific, or both (right). **C,** Venn diagram representation of the differentially expressed genes split by severity across peritoneal endometriosis (left) and percentage of severity shared genes that are non-specific, cell type-specific, tissue specific, or both (right).

**Supplementary Figure 9:**
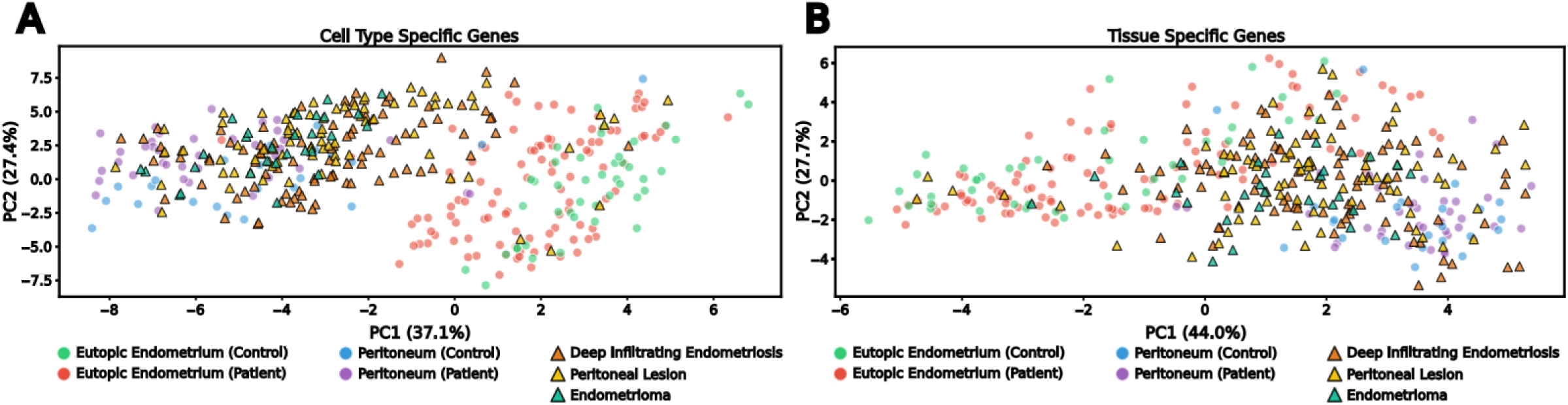
Bulk tissue microarray data can be separated by cell type- and tissue-specific gene signatures. **A-B,** PCA projections of bulk tissue microarray data using cell type-specific **(A)** genes and tissue-specific **(B)** genes.

**Supplementary Figure 10:**
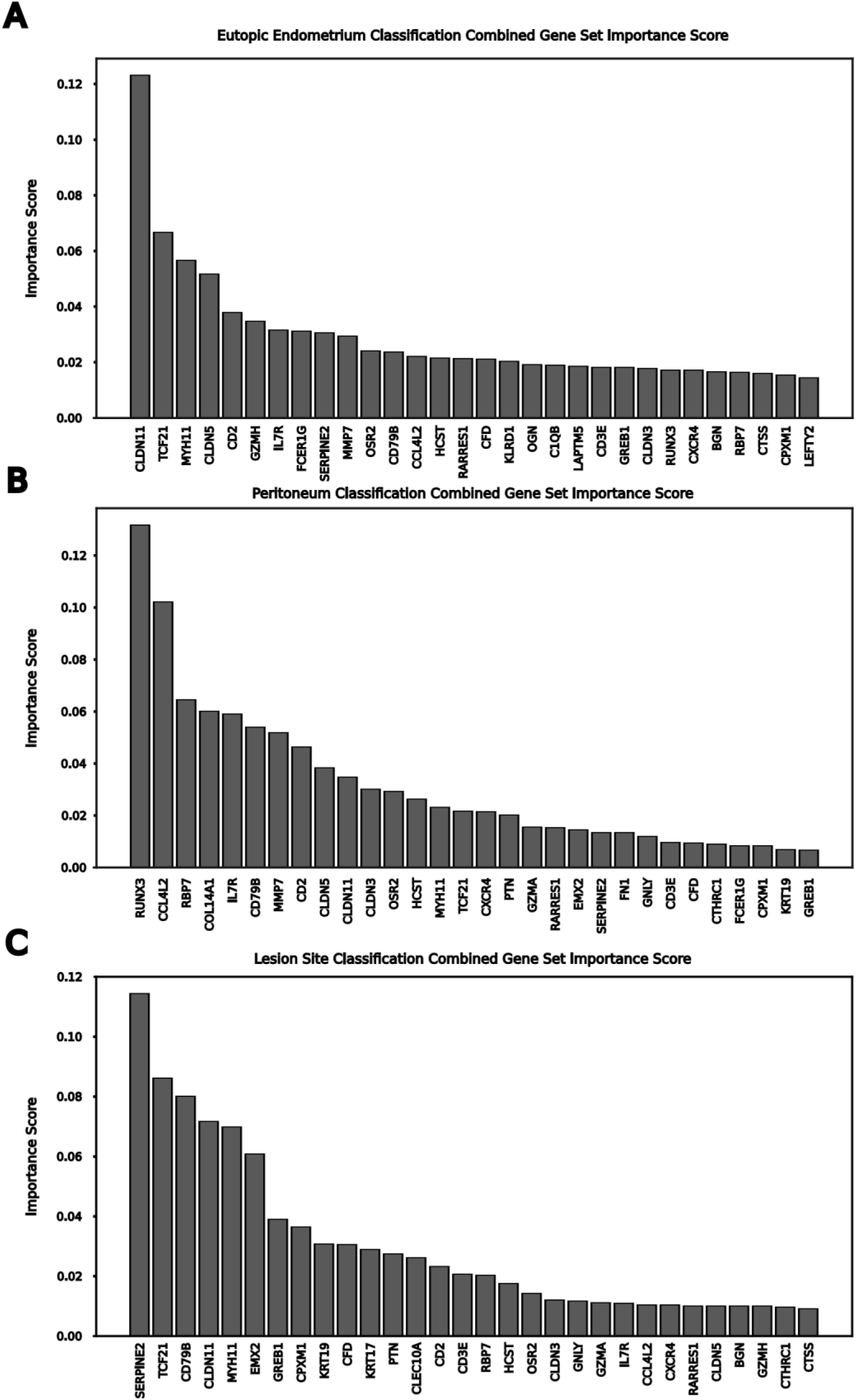
Combined cell type- and tissue-specific gene signature importances across diverse classifiers. **A,** Combined cell type- and tissue-specific gene signature classifier importances for predicting control against diseased eutopic endometrium. **B,** Combined cell type- and tissue-specific gene signature classifier importances for predicting control against diseased peritoneum. **C,** Combined cell type- and tissue-specific gene signature classifier importances for multi-class prediction across three separate lesion types (deep infiltrating endometriosis, peritoneal endometriosis, and endometrioma).

## Notes

### Competing Interest Statement

The authors have declared no competing interest.

## References

1. Zondervan, K. T., Becker, C. M. & Missmer, S. A. Endometriosis. N. Engl. J. Med. 382, 1244–1256 (2020).

2. Giudice, L. C. Endometriosis. N. Engl. J. Med. 362, 2389–2398 (2010).

3. Taylor, H. S. Reimagining Endometriosis. Med 2, 481–485 (2021).

4. Zondervan, K. T. et al. Endometriosis. Nat. Rev. Dis. Primer 4, 9 (2018).

5. American Society For Reproductive Medicine. Revised American Society for Reproductive Medicine classification of endometriosis: 1996. Fertil. Steril. 67, 817–821 (1997).

6. Nisolle, M. & Donnez, J. Peritoneal endometriosis, ovarian endometriosis, and adenomyotic nodules of the rectovaginal septum are three different entities. Fertil. Steril. 68, 585–596 (1997).

7. Bulun, S. E. et al. Endometriosis. Endocr. Rev. 40, 1048–1079 (2019).

8. Zeitoun, K. M. & Bulun, S. E. Aromatase: a key molecule in the pathophysiology of endometriosis and a therapeutic target. Fertil. Steril. 72, 961–969 (1999).

9. Vercellini, P., Viganò, P., Somigliana, E. & Fedele, L. Endometriosis: pathogenesis and treatment. Nat. Rev. Endocrinol. 10, 261–275 (2014).

10. La Marca, A., Carducci Artenisio, A., Stabile, G., Rivasi, F. & Volpe, A. Evidence for cycle-dependent expression of follicle-stimulating hormone receptor in human endometrium. Gynecol. Endocrinol. Off. J. Int. Soc. Gynecol. Endocrinol. 21, 303–306 (2005).

11. Sapkota, Y. et al. Meta-analysis identifies five novel loci associated with endometriosis highlighting key genes involved in hormone metabolism. Nat. Commun. 8, 15539 (2017).

12. Wang, W. et al. Single-cell transcriptomic atlas of the human endometrium during the menstrual cycle. Nat. Med. 26, 1644–1653 (2020).

13. Talbi, S. et al. Molecular phenotyping of human endometrium distinguishes menstrual cycle phases and underlying biological processes in normo-ovulatory women. Endocrinology 147, 1097–1121 (2006).

14. Liu, S. et al. Single-cell and spatial transcriptomic profiling revealed niche interactions sustaining growth of endometriotic lesions. Cell Genomics 5, 100737 (2025).

15. Fonseca, M. A. S. et al. Single-cell transcriptomic analysis of endometriosis. Nat. Genet. 55, 255–267 (2023).

16. Tan, Y. et al. Single-cell analysis of endometriosis reveals a coordinated transcriptional programme driving immunotolerance and angiogenesis across eutopic and ectopic tissues. Nat. Cell Biol. 24, 1306–1318 (2022).

17. Marečková, M. et al. An integrated single-cell reference atlas of the human endometrium. Nat. Genet. 56, 1925–1937 (2024).

18. Shih, A. J. et al. Single-cell analysis of menstrual endometrial tissues defines phenotypes associated with endometriosis. BMC Med. 20, 315 (2022).

19. Jones, A. S. K. et al. Cellular atlas of the human ovary using morphologically guided spatial transcriptomics and single-cell sequencing. Sci. Adv. 10, eadm7506 (2024).

20. Quake, S. R. & The Tabula Sapiens Consortium. Tabula Sapiens reveals transcription factor expression, senescence effects, and sex-specific features in cell types from 28 human organs and tissues. Preprint at 10.1101/2024.12.03.626516 (2024).

21. Crowley, G. et al. Benchmarking cell type and gene set annotation by large language models with AnnDictionary. Nat. Commun. 16, 9511 (2025).

22. Sampson, J. A. Metastatic or Embolic Endometriosis, due to the Menstrual Dissemination of Endometrial Tissue into the Venous Circulation. Am. J. Pathol. 3, 93–110.43 (1927).

23. Moore, L. et al. The mutational landscape of normal human endometrial epithelium. Nature 580, 640–646 (2020).

24. Ólafsson, S. et al. Endometriosis lesions are oligoclonal structures derived from the normal endometrium. Preprint at 10.64898/2026.02.25.708037 (2026).

25. Suda, K. et al. Different mutation profiles between epithelium and stroma in endometriosis and normal endometrium. Hum. Reprod. 34, 1899–1905 (2019).

26. Uhlén, M. et al. Proteomics. Tissue-based map of the human proteome. Science 347, 1260419 (2015).

27. Henlon, Y. et al. Single-cell analysis identifies distinct macrophage phenotypes associated with prodisease and proresolving functions in the endometriotic niche. Proc. Natl. Acad. Sci. U. S. A. 121, e2405474121 (2024).

28. Lagger, C. et al. scDiffCom: a tool for differential analysis of cell–cell interactions provides a mouse atlas of aging changes in intercellular communication. Nat. Aging 3, 1446–1461 (2023).

29. Burns, G. W. et al. Spatial transcriptomic analysis identifies epithelium-macrophage crosstalk in endometriotic lesions. iScience 28, 111790 (2025).

30. Greaves, E. et al. Estradiol is a critical mediator of macrophage-nerve cross talk in peritoneal endometriosis. Am. J. Pathol. 185, 2286–2297 (2015).

31. Ono, Y. et al. CD206+ macrophage is an accelerator of endometriotic-like lesion via promoting angiogenesis in the endometriosis mouse model. Sci. Rep. 11, 853 (2021).

32. Bacci, M. et al. Macrophages are alternatively activated in patients with endometriosis and required for growth and vascularization of lesions in a mouse model of disease. Am. J. Pathol. 175, 547–556 (2009).

33. Wu, M.-H., Hsiao, K.-Y. & Tsai, S.-J. Endometriosis and possible inflammation markers. Gynecol. Minim. Invasive Ther. 4, 61–67 (2015).

34. Dawood, M. Y. Primary dysmenorrhea: advances in pathogenesis and management. Obstet. Gynecol. 108, 428–441 (2006).

35. Stratton, P. & Berkley, K. J. Chronic pelvic pain and endometriosis: translational evidence of the relationship and implications. Hum. Reprod. Update 17, 327–346 (2011).

36. ACOG Committee Opinion No. 760: Dysmenorrhea and Endometriosis in the Adolescent. Obstet. Gynecol. 132, e249–e258 (2018).

37. Begum, I. A. The connection between endometriosis and secondary dysmenorrhea. J. Reprod. Immunol. 168, 104425 (2025).

38. Deuster, P. A., Dolev, E., Bernier, L. L. & Trostmann, U. H. Magnesium and zinc status during the menstrual cycle. Am. J. Obstet. Gynecol. 157, 964–968 (1987).

39. Bruner, K. L., Eisenberg, E., Gorstein, F. & Osteen, K. G. Progesterone and transforming growth factor-â coordinately regulate suppression of endometrial matrix metalloproteinases in a model of experimental endometriosis. Steroids 64, 648–653 (1999).

40. Obeagu, E. I. & Obeagu, G. U. From menses to ovulation: the immunomodulatory role of monocyte cytokines in menstrual cycle physiology and reproductive health - a narrative review. Ann. Med. Surg. 87, 8453–8459 (2025).

41. Wira, C. R., Rodriguez-Garcia, M. & Patel, M. V. The role of sex hormones in immune protection of the female reproductive tract. Nat. Rev. Immunol. 15, 217–230 (2015).

42. Attia, G. R. et al. Progesterone receptor isoform A but not B is expressed in endometriosis. J. Clin. Endocrinol. Metab. 85, 2897–2902 (2000).

43. Burney, R. O. et al. Gene expression analysis of endometrium reveals progesterone resistance and candidate susceptibility genes in women with endometriosis. Endocrinology 148, 3814–3826 (2007).

44. Vissers, G., Giacomozzi, M., Verdurmen, W., Peek, R. & Nap, A. The role of fibrosis in endometriosis: a systematic review. Hum. Reprod. Update 30, 706–750 (2024).

45. Gong, H. et al. The regulation of ovary and conceptus on the uterine natural killer cells during early pregnancy. Reprod. Biol. Endocrinol. 15, 73 (2017).

46. Hogg, C. et al. Macrophages inhibit and enhance endometriosis depending on their origin. Proc. Natl. Acad. Sci. 118, e2013776118 (2021).

47. Rocha, A. L. L., Reis, F. M. & Taylor, R. N. Angiogenesis and endometriosis. Obstet. Gynecol. Int. 2013, 859619 (2013).

48. Reichelt, U. et al. High lymph vessel density and expression of lymphatic growth factors in peritoneal endometriosis. Reprod. Sci. 19, 876–882 (2012).

49. Zhang, Q., Duan, J., Olson, M., Fazleabas, A. & Guo, S.-W. Cellular Changes Consistent With Epithelial-Mesenchymal Transition and Fibroblast-to-Myofibroblast Transdifferentiation in the Progression of Experimental Endometriosis in Baboons. Reprod. Sci. 23, 1409–1421 (2016).

50. Pušić, M. N. et al. Growth Arrest-Specific Protein 6 Is Elevated in Endometriosis but Shows Poor Diagnostic Performance. Int. J. Mol. Sci. 26, 8348 (2025).

51. Zhai, X. et al. Gas6/AXL pathway: immunological landscape and therapeutic potential. Front. Oncol. 13, 1121130 (2023).

52. Klemmt, P. A. B., Carver, J. G., Kennedy, S. H., Koninckx, P. R. & Mardon, H. J. Stromal cells from endometriotic lesions and endometrium from women with endometriosis have reduced decidualization capacity. Fertil. Steril. 85, 564–572 (2006).

53. Wang, X. et al. The role of insulin-like growth factor binding proteins in TGF-β1-induced fibroblast-myofibroblast transition during endometriosis fibrosis. Cell. Signal. 140, 112362 (2026).

54. Giudice, L. C., Dsupin, B. A., Gargosky, S. E., Rosenfeld, R. G. & Irwin, J. C. The insulin-like growth factor system in human peritoneal fluid: its effects on endometrial stromal cells and its potential relevance to endometriosis. J. Clin. Endocrinol. Metab. 79, 1284– 1293 (1994).

55. Clower, L., Fleshman, T., Geldenhuys, W. J. & Santanam, N. Targeting Oxidative Stress Involved in Endometriosis and Its Pain. Biomolecules 12, 1055 (2022).

56. Harrington, D. J. et al. Tenascin is differentially expressed in endometrium and endometriosis. J. Pathol. 187, 242–248 (1999).

57. Ma, R. et al. Identification of key genes associated with endometriosis and endometrial cancer by bioinformatics analysis. Front. Oncol. 14, 1387860 (2024).

58. Yotova, I. et al. LINC01638 promotes epithelial-to-mesenchymal transition in endometriosis epithelial cells by up-regulating RHOB via HDAC1 suppression. Reprod. Biomed. Online 51, 104942 (2025).

59. Ganieva, U. et al. Involvement of Transcription Factor 21 in the Pathogenesis of Fibrosis in Endometriosis. Am. J. Pathol. 190, 145–157 (2020).

60. Lee, M.-Y. et al. Role of interleukin-32 in the pathogenesis of endometriosis: in vitro, human and transgenic mouse data. Hum. Reprod. 33, 807–816 (2018).

61. Rahmioglu, N. et al. Genetic variants underlying risk of endometriosis: insights from meta-analysis of eight genome-wide association and replication datasets. Hum. Reprod. Update 20, 702–716 (2014).

62. Vorperian, S. K. et al. Cell types of origin of the cell-free transcriptome. Nat. Biotechnol. 40, 855–861 (2022).

63. Moufarrej, M. N. et al. Early prediction of preeclampsia in pregnancy with cell-free RNA. Nature 602, 689–694 (2022).

64. Gabriel, M. et al. A relational database to identify differentially expressed genes in the endometrium and endometriosis lesions. Sci. Data 7, 284 (2020).

65. Li, Q. et al. Identification of Candidate Gene Signatures and Regulatory Networks in Endometriosis and its Related Infertility by Integrated Analysis. Reprod. Sci. 29, 411– 426 (2022).

66. Donnez, J. & Cacciottola, L. Endometriosis: An Inflammatory Disease That Requires New Therapeutic Options. Int. J. Mol. Sci. 23, 1518 (2022).

67. Lousse, J.-C. et al. Peritoneal endometriosis is an inflammatory disease. Front. Biosci. 4, 23–40 (2012).

68. Ferrero, S., Vellone, V. G. & Barra, F. Pathophysiology of pain in patients with peritoneal endometriosis. Ann. Transl. Med. 7, S8 (2019).

69. Nisenblat, V. et al. Blood biomarkers for the non-invasive diagnosis of endometriosis. Cochrane Database Syst. Rev. 2016, CD012179 (2016).

70. Halpern, K. B. et al. Single-cell spatial reconstruction reveals global division of labour in the mammalian liver. Nature 542, 352–356 (2017).

71. Moor, A. E. et al. Spatial Reconstruction of Single Enterocytes Uncovers Broad Zonation along the Intestinal Villus Axis. Cell 175, 1156–1167.e15 (2018).

72. Suda, K. et al. Clonal Expansion and Diversification of Cancer-Associated Mutations in Endometriosis and Normal Endometrium. Cell Rep. 24, 1777–1789 (2018).

73. Anglesio, M. S. et al. Cancer-Associated Mutations in Endometriosis without Cancer. N. Engl. J. Med. 376, 1835–1848 (2017).

74. Murray, P. J. & Wynn, T. A. Protective and pathogenic functions of macrophage subsets. Nat. Rev. Immunol. 11, 723–737 (2011).

75. Donnez, J. et al. Peritoneal endometriosis and “endometriotic” nodules of the rectovaginal septum are two different entities. Fertil. Steril. 66, 362–368 (1996).

76. Lavin, Y. et al. Tissue-resident macrophage enhancer landscapes are shaped by the local microenvironment. Cell 159, 1312–1326 (2014).

77. Gautier, E. L. et al. Gene-expression profiles and transcriptional regulatory pathways that underlie the identity and diversity of mouse tissue macrophages. Nat. Immunol. 13, 1118–1128 (2012).

78. Koh, W. et al. Noninvasive in vivo monitoring of tissue-specific global gene expression in humans. Proc. Natl. Acad. Sci. 111, 7361–7366 (2014).

79. Larson, M. H. et al. A comprehensive characterization of the cell-free transcriptome reveals tissue- and subtype-specific biomarkers for cancer detection. Nat. Commun. 12, 2357 (2021).

80. Warren, L. A. et al. Analysis of menstrual effluent: diagnostic potential for endometriosis. Mol. Med. 24, 1 (2018).

81. 81. Wilson, T. R. et al. Analysis of menstrual effluent uncovers endometriosis-specific cell populations and impaired cellular pathway processes. Preprint at 10.1101/2025.08.21.671582 (2025).

82. Garcia-Alonso, L. et al. Mapping the temporal and spatial dynamics of the human endometrium in vivo and in vitro. Nat. Genet. 53, 1698–1711 (2021).

83. Lai, Z.-Z. et al. Single-cell transcriptome profiling of the human endometrium of patients with recurrent implantation failure. Theranostics 12, 6527–6547 (2022).

84. Huang, X. et al. Single-cell transcriptome analysis reveals endometrial immune microenvironment in minimal/mild endometriosis. Clin. Exp. Immunol. 212, 285–295 (2023).

85. Korsunsky, I. et al. Fast, sensitive and accurate integration of single-cell data with Harmony. Nat. Methods 16, 1289–1296 (2019).

86. Muyas, F. et al. De novo detection of somatic mutations in high-throughput single-cell profiling data sets. Nat. Biotechnol. 42, 758–767 (2024).

87. Yanai, I. et al. Genome-wide midrange transcription profiles reveal expression level relationships in human tissue specification. Bioinformatics 21, 650–659 (2005).

88. Finak, G. et al. MAST: a flexible statistical framework for assessing transcriptional changes and characterizing heterogeneity in single-cell RNA sequencing data. Genome Biol. 16, 278 (2015).

